# Genetic analysis of *Mycobacterium abscessus* reveals genomic diversity linked to global plasmid distribution

**DOI:** 10.1101/2024.06.11.597493

**Authors:** Kensuke Ohse, Atsushi Yoshida, Keisuke Kamada, Hironobu Kitazawa, Yusuke Ito, Takayo Shoji, Kenichiro Watanabe, Hiroshi Koganemaru, Ken Kikuchi, Masashi Toyoda

## Abstract

Genetic epidemiological analysis of mobile genetic elements, such as plasmids, has rarely been carried out in *Mycobacterium abscessus,* regardless of its usefulness in speculating the past contact between bacteria. In this study, through whole genome sequencing analysis of clinical isolates of *M. abscessus* sequentially collected from the same patient, we identified a single nucleotide variant that could cause antibiotic resistance, and three plasmids (pMAB625-1, pMAB625-2 and pMAB625-3) not found in the type strain. We investigated the distribution of plasmids previously identified in *M. abscessus,* including pMAB625 plasmids, in 462 clinical isolates worldwide *in silico*. pMAB625-3 was detected in the largest number of isolates. Furthermore, phylogenetic tree analysis revealed that these plasmids transferred beyond regions and subspecies and acquired unique mutations. These results indicate that transmission of plasmids increases the genomic diversity in *M. abscessus,* and plasmid epidemiology is useful for estimating the past contact between bacteria.

## INTRODUCTION

In recent years, the threat of antimicrobial-resistant bacteria has been increasing worldwide, and they could be responsible for 1.91 million deaths per year by 2050^1^. Hence, research into the mechanisms and routes of transmission is becoming increasingly important to develop effective countermeasures against antimicrobial-resistant bacteria. The global incidence of non-tuberculous mycobacterial infections has been on the rise, presenting a significant challenge due to their increasing prevalence and innate drug resistance, particularly in *Mycobacterium abscessus* infections^2^. *M. abscessus* is not limited to pulmonary infections but also serves as the causative agent for various extrapulmonary infections, including bloodstream, skin, soft tissue, surgical site, and peritoneal dialysis-related infections^3^. Furthermore, it demonstrates resistance to a wide range of antimicrobial agents.

The route of infection of *M. abscessus* remains controversial. It has been assumed that *M. abscessus* is transmitted from the environment, such as soil and water systems, and person-to-person transmission does not occur. However, a few cases of suspected person-to-person transmission have been reported^4–6^. In addition, the existence of “dominant clones”, which were isolated in almost all regions and phylogenetically close^6,7^, makes the source and transmission routes of the bacteria mysterious^8^.

Whole genome sequencing (WGS) analysis and phylogenetic tree analysis of the core genome, which is the genomic region shared by all isolates, are often conducted to predict the origin and transmission route of the isolates^9^. However, the accessory genome, which is shared by a certain percentage but not all of the isolates such as plasmids, prophage, and genomic islands, is made up of approximately 30% of the typical number of all genes of *M. abscessus*^10^. Therefore, the analysis of the pan-genome, which includes both the core genome and accessory genome, was performed to compare the genetic relation of all gene sets of *M. abscessus*^8,11^. Although the pan-genome analysis has the advantage of comparing whole bacterial genomes, it has the disadvantage of analyzing accessory genomes as gene units, resulting in the loss of information on the units of mobile genetic elements. Therefore, epidemiological analysis of mobile genetic element units, such as individual plasmids, may more accurately predict the origin and route of infection of bacteria.

In this study, we performed the WGS analysis of the clinical isolates that acquired elevated minimum inhibitory concentrations (MICs) for carbapenem antibiotics in a case of disseminated infection caused by *M. abscessus* subsp. *abscessus* (ABS) during antimicrobial therapy. We detected one mutation potentially responsible for antimicrobial resistance (AMR) and identified three plasmids (pMAB625-1, pMAB625-2 and pMAB625-3) that type strain did not harbor. Therefore, we analyzed the distribution of plasmids identified in *M. abscessus* before, including pMAB625 plasmids and found that pMAB625-3 was the most widely spreading plasmid among the clinical isolates of *M. abscessus* worldwide. Thus, we investigated whether predicting the past contact between isolates is possible by analyzing single nucleotide variants (SNVs) and structural variants (SVs) of the plasmids.

## RESULTS

### The clinical isolates had decreased susceptibility to carbapenem antibiotics compared to the type strain

We sequentially isolated five ABS strains from one patient who had osteomyelitis over six months (Table 1). We first compared the growth rate and antimicrobial susceptibility of these clinical isolates with the type strain (ATCC 19977). The growth rates of Isolate57625 and Isolate57626 were significantly slower than the type strain (Figure S1). Antimicrobial susceptibility testing showed that the MIC of Isolate57626 against IPM, which binds the penicillin-binding proteins (PBPs) and inhibits the bacterial cell wall synthesis, was higher than that of the type strain (Table 1). The MICs of Isolate57625 and Isolate57626 against meropenem (MEPM), which has the same mechanism of action as IPM and improved stability, were higher than that of the type strain (Table 1). However, there were no differences in MIC between ATCC 19977 and two isolates against the other antibiotics. These results indicated the decreased susceptibility of the isolates to carbapenem antibiotics compared to the type strain.

**Table 1.** The isolation time point of the clinical isolates and antimicrobial susceptibility of the type strain and the isolates.

| Isolates | Source | Period from the first isolation (month) | MIC<sup>a</sup> (µg/mL) |  |  |  |  |  |  |  |  |  |  |  |  |  |  |
| --- | --- | --- | --- | --- | --- | --- | --- | --- | --- | --- | --- | --- | --- | --- | --- | --- | --- |
|  |  |  | AMK | TOb | IPM | FRPM | LVFX | MFLX | AZM | CAM 4day | CAM 14day | ST | DOXY | MEPM | LZD | CLF | STFX |
| ATCC 19977 | - | - | 8 | 8 | 16 | >64 | 32 | >8 | 64 | 2 | >64 | >152/8 | >16 | 64 | 32 | 1 | 2 |
| Isolate57625 | catheter | 0 | 8 | 8 | 16<sup>b</sup> | >64 | 16 | >8 | 64 | 2 | >64 | >152/8 | >16 | >64 | 32 | 1 | 2 |
| Isolate57629 | catheter | 4 | - | - | - | - | - | - | - | - | - | - | - | - | - | - | - |
| Isolate57630 | bone biopsy | 4 | - | - | - | - | - | - | - | - | - | - | - | - | - | - | - |
| Isolate57596 | heel | 6 | - | - | - | - | - | - | - | - | - | - | - | - | - | - | - |
| Isolate57626 | malleolus | 6 | 8 | 8 | >64<sup>c</sup> | >64 | 32 | >8 | 64 | 2 | >64 | >152/8 | >16 | >64 | 32 | 1 | 2 |

<sup>a</sup>MICs were determined as the minimum concentration at which the bacteria did not grow. No antimicrobial susceptibility testing was conducted on the isolates other than Isolate57625 and Isolate57626, representing hyphens in the table. The test was repeated three times and all MICs were confirmed to be the same between the experiments with the exception of imipenem for the isolates. AMK, Amikacin; TOB, Tobramycin; IPM, Imipenem; FRPM, Faropenem; LVFX, Levofloxacin; MFLX, Moxifloxacin; AZM, Azithromycin; CAM, Clarithromycin; ST, Trimethoprim/sulfamethoxazole; DOXY, Doxycycline; MEPM, Meropenem; LN2, Linezolid; CLF, Clofazimine; STFX, Sitafloxacin
<sup>b</sup>For Isolate57625, the MICs against IPM in the three experiments were 64, 16, and 16 µg/mL, respectively.
<sup>c</sup>For Isolate57626, the MICs against IPM in the three experimenst were >64, 16, and >64 µg/mL, respectively.

### WGS of the clinical isolates

To explore the factors determining the difference in growth rate and antimicrobial susceptibility between the type strain and the clinical isolates, we performed WGS using short-read sequencer and Nanopore long-read sequencer. We analyzed the *de novo* assembled chromosome sequences and found that two large insertions (13 kbp and 8 kbp) and a deletion (16 kbp) existed in the sequence of Isolate57625 (Figure S2A, B). The coding sequences of phage integrase and transposase were present near the three large SVs (Figure S3), indicating that these regions were prophage and transposon. We also analyzed the SNVs in the chromosome (Table S3), however, we could not identify the cause of the phenotypic differences. Phylogenetic analysis showed that the isolates were genetically close to the type strain (Figure S2C). Furthermore, to analyze the difference in the chromosome sequence between five clinical isolates, we analyzed SVs and SNVs between the isolates (Figure S4). This analysis suggested one mutation located at the putative *PbpA*, which was associated with the synthesis of the bacterial cell wall^12^ and the target of carbapenem antibiotics^13^, may be responsible for the reduced susceptibility of Isolate57626 to IPM compared to Isolate57625.

### The isolates had three plasmids that encode various virulence factors

WGS analysis also revealed that the clinical isolates have three plasmids not found in the type strain (named pMAB625-1, pMAB625-2 and pMAB625-3, respectively) (Table 2). Thus, we next focused on the pMAB625 plasmids and compared the pMAB625 plasmid sequences with other plasmids registered to PLSDB^14^ previously (Table 3). A taxonomic keyword search for “*abscessus*” returned 20 plasmids. To investigate the homology between these plasmids, we calculated ANI between them and pMAB625. No plasmid had a high ANI for pMAB625-1, however, the ANI between pMAB625-2 and pGD21-2 was 97.4% with some SVs. Furthermore, the ANI for pMAB625-3 against pGD69A-1 and pGD42-1 was 100%. These results indicated that pMAB625-2 and pMAB625-3 were retained in the other clinical isolates of *M. abscessus.* Next, we performed qPCR using primers specific to each plasmid to verify the copy numbers of pMAB625 plasmids in the clinical isolates. This analysis showed that the copy number of pMAB625-1 and pMAB625-2 was less than one per bacterium, and that of pMAB625-3 was more than five per bacterium in the isolates (Figure S5). Therefore, the clinical isolates were considered to be a heterogeneous population in terms of plasmid possession.

**Table 2.** Genome assembly and the annotation statics of the Isolate57625.

| Contig | Form | Length(bp) | GC contents (%) | Number of CDS | Number of rRNA | Number of tRNA | Number of tmRNA |
| --- | --- | --- | --- | --- | --- | --- | --- |
| Chromosome | circular | 5,072,346 | 64.1 | 5,034 | 3 | 48 | 1 |
| pMAB625-1 | linear | 202,398 | 64.7 | 247 | 0 | 29 | 0 |
| pMAB625-2 | linear | 142,689 | 62.2 | 171 | 0 | 0 | 0 |
| pMAB625-3 | circular | 24,988 | 65.6 | 30 | 0 | 0 | 0 |

**Table 3.** Homology search result between plasmids identified in *M. abscessus*.

| Plasmid | Accession no. | Topology | Length (bp) | Homologous plasmid (ANI(>99)) | Homologous plasmid (97< ANI(<99) ) | Use for "Reference" |
| --- | --- | --- | --- | --- | --- | --- |
| pMAB625-1 | AP028614.1 | linear | 202,398 | - | - | TRUE |
| pMAB625-2 | AP028615.1 | linear | 142,689 | - | pGD21-2<sup>b</sup> | TRUE |
| pMAB625-3 | AP028616.1 | circular | 24,988 | pGD69A-1, pGD42-1 | - | TRUE |
| pMAB23 | NC_010394.1 | circular | 23,319 | pMabS_GZ002<sup>a</sup>, FDAARGOS_1604_plasmid | - | TRUE |
| pMabS_GZ002 | NZ_CP034180.1 | linear | 15,203 | pMAB23, FDAARGOS_1604_plasmid | - | TRUE |
| pGD19 | NZ_CP063329.1 | circular | 18,605 | - | - | TRUE |
| pGD21-1 | NZ_CP065285.1 | circular | 112,633 | - | - | TRUE |
| pGD21-2 | NZ_CP065286.1 | linear | 155,609 | - | pMAB625-2<sup>b</sup> | TRUE |
| pGD22-1 | NZ_CP063325.1 | circular | 19,694 | - | - | TRUE |
| pGD22-2 | NZ_CP063326.1 | circular | 18,117 | pGD42-2<sup>a</sup> | - | TRUE |
| pGD25-1 | NZ_CP063321.1 | circular | 31,413 | - | - | TRUE |
| pGD25-2 | NZ_CP063322.1 | circular | 27,424 | - | - | TRUE |
| pGD25-3 | NZ_CP063323.1 | circular | 23,599 | - | - | TRUE |
| pGD42-1 | NZ_CP065281.1 | circular | 24,993 | pMAB625-3, pGD69A-1 | - | FALSE |
| pGD42-2 | NZ_CP065282.1 | circular | 9,547 | pGD22-2 | pGD69A-2 | TRUE |
| pGD69A-1 | NZ_CP065270.1 | circular | 25,000 | pMAB625-3, pGD42-1 | - | FALSE |
| pGD69A-2 | NZ_CP065271.1 | circular | 18,611 | - | pGD42-2<sup>a</sup> | TRUE |
| pJCM30620_1 | NZ_AP022622.1 | linear | 119,389 | 50594_plasmid2<sup>a</sup> | - | TRUE |
| pJCM30620_2 | NZ_AP022623.1 | linear | 38,763 | 50594_plasmid1<sup>b</sup> | - | TRUE |
| BRA100 | NC_017908.2 | circular | 56,265 | - | - | TRUE |
| FDAARGOS_1<br>604_plasmid | NZ_CP085920.1 | circular | 23,319 | pMAB23,<br>pMabS_GZ002<sup>a</sup> | - | FALSE |
| 50594_plasmid<br>d1 | NC_021278.1 | circular | 172,814 | pJCM30620_2<sup>a</sup> | - | TRUE |
| 50594_plasmid<br>d2 | NC_021279.1 | circular | 97,240 | pJCM30620_1<sup>b</sup> | - | TRUE |

<sup>a</sup>One Structural Variant (SV) exists in the sequence.
<sup>b</sup>More than two SVs exist in the sequence.

We then investigated the function of the genes encoded on pMAB625 plasmids using homology search. Plasmids encode various factors that influence host characteristics. Therefore, we listed the lost and acquired genes in Isolate57625 compared to the type strain to investigate the factors affecting antimicrobial susceptibility (Table S4). Of note, 480 candidate open reading frames (ORFs) were acquired and 37 candidate ORFs were lost in Isolate57625 compared to the type strain. We then searched the AMR genes from several databases using Blast+ based on the amino acid sequences of the acquired candidate ORFs. This analysis showed that the coding sequence (CDS) named Isolate57625_54550 had a similarity to *qacA*, a subunit of the qac multidrug efflux pump^15^, (identity, 31 %; *e*-value, 1.69e-10; coverage, 16.1%) and that the CDS named Isolate57625_54800 had a similarity to *MexL*, a specific repressor of *MexJK* multidrug efflux system^16^ (identity, 33%; *e*-value, 5.54e-4; coverage, 34.4%). However, the identity and coverage were not sufficiently high.

To gain further insight into the effect of gene acquisition, we searched the homology of the acquired candidate ORFs using the mycobacterial gene database of Mycobrowser^17^. This analysis detected 37 mycobacterial proteins with amino acid homology to candidate ORFs of the isolates (identity > 30 %, *e*-value < 1e-50) (Table S5). Notably, genes associated with the ESX secretion system were highly enriched (10/37, 27%). ESX secretion systems are encoded in various bacteria on their chromosomes and plasmids. It has been reported that the ESX secretion system (also known as the type VII secretion system) is involved in bacterial virulence through several mechanisms, including evasion of the host immune system^18,19^, and biofilm formation^20^. We further searched whether other ORFs of the components of the ESX secretion system were encoded near the loci of the detected ORFs of the components of the ESX secretion system using blast+. We detected several components of the ESX secretion system in pMAB625-1 and pMAB625-2 (Figure 1A). Core genes that construct membrane pores (*eccB, eccC, eccD, mycP,* and *eccE*) were present in these gene clusters. The type strain and clinical isolates have ESX-3 and ESX-4 loci on their chromosomes; therefore, the isolates acquired two additional ESX loci by the plasmid acquisition. To predict the function of these genes, we performed phylogenetic tree analysis using known amino acid sequences of the ESX secretion system components in chromosomes and plasmids. It was previously reported that plasmid-encoded ESX loci can be classified into 4 major clusters^21^. Importantly, our analysis revealed that plasmid-encoded ESX loci were classified into 5 major clusters (ESX-P cluster 1-5) (Figure 1B). Notably, the newly classified ESX-P cluster 5 contained the ESX loci of pMAB625-1 and pMAB625-2. As shown in the results of the plasmid homology search (Table 3), the ESX loci of pGD21-2 and pMAB625-2 were genetically close. The ESX loci of pMAB625-1 were genetically close to that of pMFLV01, which was harboured by *Mycobacterium gilvum* PYR GCK. It was previously reported that pMFLV01 was enriched in biofilm formed in household water purifiers by metagenomic analysis^22^. Biofilm formation is one of the drug resistance mechanisms in mycobacterium and acts as a barrier against antibiotics. We also detected the toxin-antitoxin (TA) system components, YefM/YoeB and VapB5/VapC5 in pMAB625-1. The TA system is present in various bacteria and archaea and encoded on their chromosomes and plasmids. It contributes to plasmid persistence, inhibiting bacteriophage propagation, and increases antimicrobial tolerance by inducing dormancy^23^. These results indicated that the clinical isolates acquired factors that could contribute to bacterial virulence through plasmid transfer.

**Figure 1.**
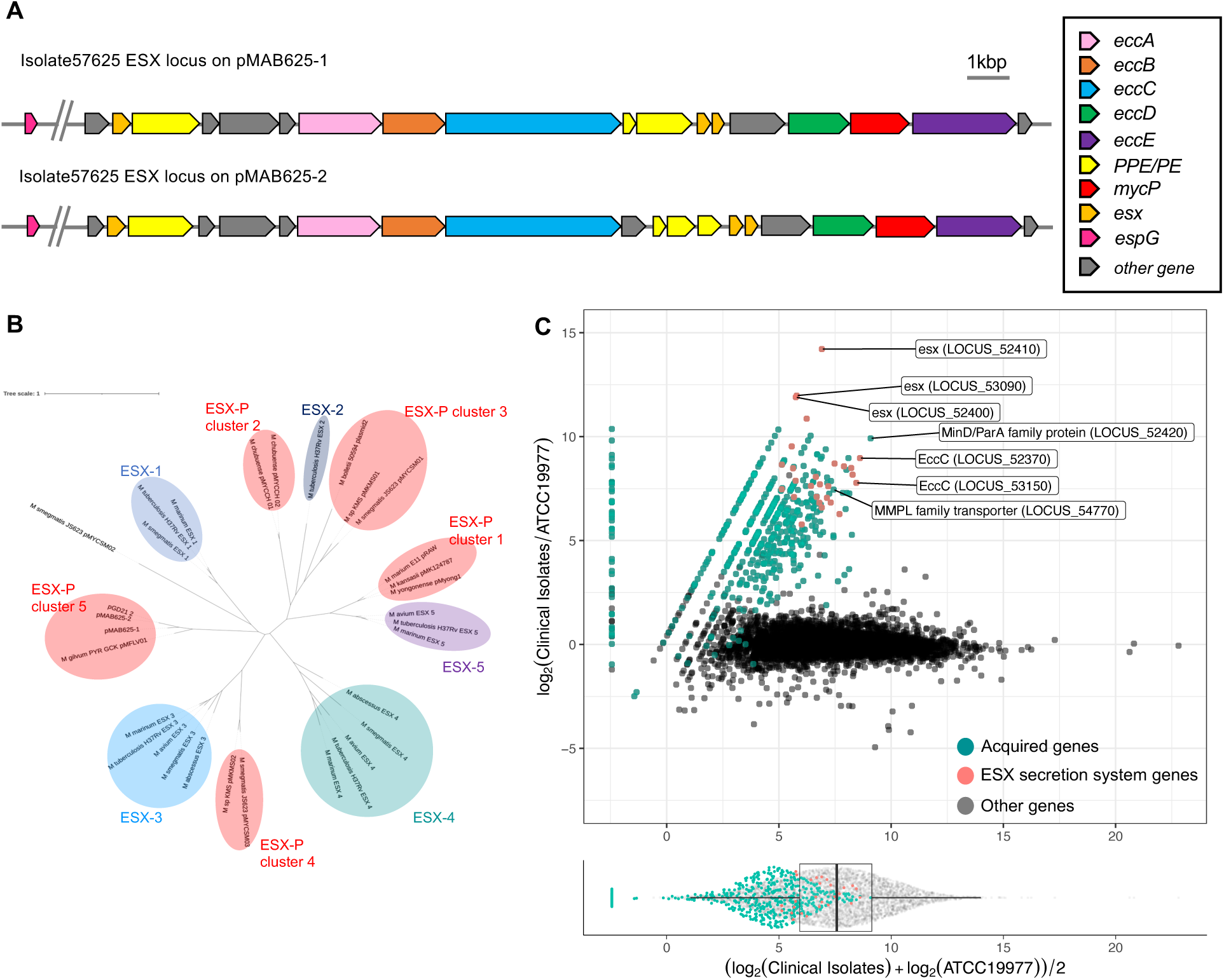
Phylogenetic and gene expression analysis of the genes related to the ESX secretion system (A) The loci of the ESX secretion system are encoded on pMAB625-1 and pMAB625-2. CDSs of the ESX secretion system were searched using Blast+. The core components that construct membrane pore (*eccB, eccC, eccD, mycP,* and *eccE*) are contained in this operon. (B) Phylogenetic tree analysis of various ESX loci encoded on chromosomes and plasmids. The amino acid sequence of the genes that construct the membrane pore were concatenated and the phylogenetic tree was created by NGphylogeny.fr^24^ using the PhyML+SMS inference method. The classification of each cluster was followed as previously reported^21^. (C) RNA-seq analysis of the type strain and the isolates. The x-axis represents the mean log expression levels, and the y-axis of the top plot represents the log fold change. The bottom dot plot and box plot represent the distribution of the expression levels of each gene. The green dots represent the data of genes that the isolates acquired by SVs and plasmid acquisition. The red dots represent the data of ESX secretion system genes encoded on the plasmids.

### Gene expression analysis on the pMAB625 plasmids in the clinical isolates

We found that the various genes associated with bacterial virulence were encoded on the pMAB625 plasmid. However, it was unclear whether genes encoded on these plasmids were expressed in the isolates and to what extent gene expression on the plasmids influenced the expression of genes on the chromosome. To address this question, we analyzed the global gene expression in the type strain and the isolates using RNA-seq analysis (Figure 1C and Table S6). The expression levels of the genes on plasmids were low to moderate (below the 75th percentile of the distribution of expression levels of all genes that were expressed in the isolates and the type strain). Notably, the expression levels of the most genes (19/27 genes) associated with the ESX secretion system, which were encoded on the pMAB625-1 and pMAB625-2, were moderate (below the 75th percentile and above the 25th percentile) (Figure 1C). In addition, the genes associated with the TA system and the mycobacterial membrane protein large (MMPL) family transporter, which were also encoded on the pMAB625-1 and pMAB625-3, respectively, were expressed at moderate levels. These results indicated that the numerous genes associated with bacterial virulence encoded on the plasmids were expressed at moderate levels in the clinical isolates, despite the low number of copies per bacterium for the pMAB625-1 and pMAB625-2. Next, in the clinical isolates, 393 genes were highly expressed and 29 genes were low expressed compared to the type strain. Of these differentially expressed genes, 87.2% of genes (368 highly expressed genes) were acquired genes by SVs or plasmids, and 12.8% of the genes (25 highly expressed genes and 29 low expressed genes) were present on the chromosomes of both the isolates and the type strain (Figure 1C). These results indicated that the gene expression on the plasmid could directly or indirectly affect the expression on the chromosome but to a lesser extent.

### pMAB625 plasmids were globally distributed in the clinical isolates of M. abscessus

So far, we have found that the isolates carried three plasmids that the type strain did not, that these plasmids encoded the genes that could contribute to bacterial virulence, and that these genes also expressed in the isolates. However, we had analyzed the strains sequentially isolated from only one patient and it was uncertain whether the plasmid acquisition occurred extensively in clinical isolates of mycobacterium. We therefore investigated whether other clinical isolates of mycobacterium carry the pMAB625 plasmids. We first performed PCR using the genomic DNA of the clinical isolates collected in Japan^25^ with primers specific to each pMAB625 plasmid. We detected pMAB625-3 in two isolates other than Isolate57625, although we did not detect pMAB625-1 and pMAB625-2 in any isolates other than Isolate57625 (Figure 2A). Both the isolates in which pMAB625-3 was detected were *M. abscessus*. We then verified the plasmid possession by mapping the read data from the short-read sequencer to the pMAB625 plasmid sequences. We used the read data of *M. abscessus* clinical isolates in Japan^7^ obtained from the NCBI database in addition to our original data. Out of 125 clinical isolates, pMAB625-2 was detected in four isolates (3.2%) and pMAB625-3 was detected in 16 isolates (12.8%) (Figure 2B). These isolates were collected from the four prefectures in Japan (Tokyo, Shizuoka, Aomori, and Okinawa) (Figure 2C). We also mapped the read data of the other mycobacterium species (three of *Mycobacterium iranicum*, 20 of *Mycobacterium fortuitum*, and 25 of *Mycobacterium chelonae*), however, no pMAB625 plasmid was detected in these isolates (data not shown).

**Figure 2.**
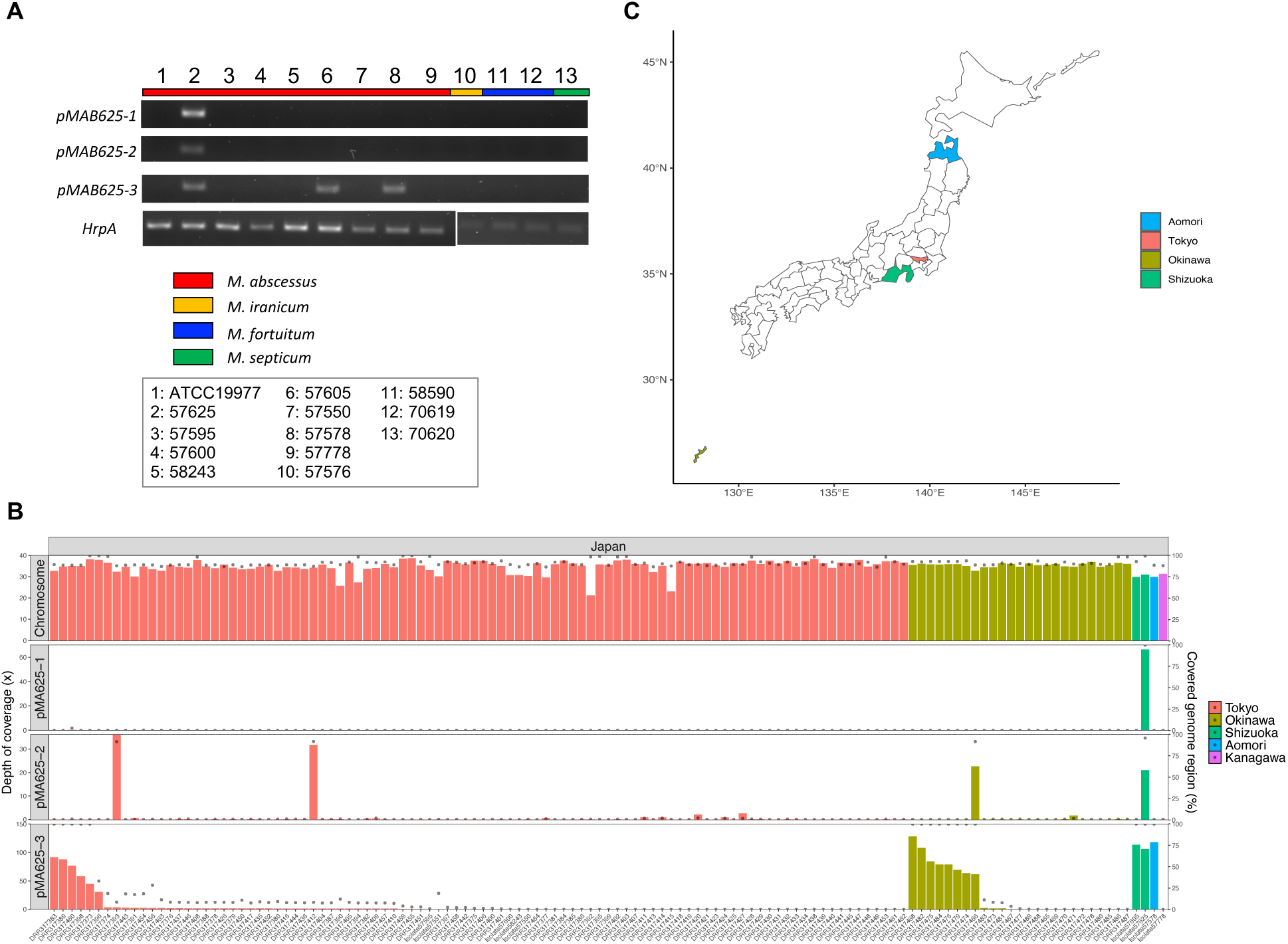
The distribution of pMAB625 plasmid in the clinical isolates in Japan (A) PCR analysis of the possession of pMAB625 plasmid in the clinical isolates which our group isolated in Japan. PCR was performed using the primers specific to each plasmid and *HrpA*. The color bar represents the species of each isolate. The primer sets of *HrpA* were different depending on the species. (B) Mapping the read data of the clinical isolates in Japan to Isolate57625 genome sequence. The bar plot represents the depth of coverage to chromosome and each plasmid and the dot plot represents the percentage of covered genome region. (C) Geographical map of Japan. The prefectures in which the pMAB625 plasmids were detected are represented by color.

Moreover, to investigate whether clinical isolates in other areas of East Asia possess pMAB625 plasmids, we mapped the available read data about the clinical isolates from Taiwan^7^ and China^26^. pMAB625-2 was detected in the isolates from China (2/69, 2.9%) and pMAB625-3 was detected in the isolates from Taiwan (8/98, 8.2%), and China (5/69, 7.2%) (Figure S6). Two plasmids were detected in ABS and *M. abscessus* subsp. *massiliense* (MAS) but not in *M. abscessus* subsp. *bolletii* (BOL). In addition, we analyzed the clinical isolates in Spain, the UK, and the US to determine whether these isolates carried the pMAB625 plasmids (Table 4). pMAB625-2 plasmid was detected in isolates from the UK (2/30, 6.7%), and the US (2/110, 1.8%). pMAB625-3 plasmid was detected in the isolates from Spain (9/30, 30%), the UK (1/30, 3.3%), and the US (14/110, 12.7%). Overall, pMAB625-1 was detected in one isolate (1/462, 0.2%), pMAB625-2 was detected in 10 isolates (10/462, 2.2%), and pMAB625-3 was detected in 53 isolates (53/462, 11.5%).

**Table 4.** pMAB625 and known plasmids distribution in the clinical isolates worldwide.

| Number (%) of isolates retaining plasmid |  |  |  |  |  |  |  |  |
| --- | --- | --- | --- | --- | --- | --- | --- | --- |
| Region | Unknown<br>(Type strain) | Japan | China | Taiwan | US | Spain | UK | total<sup>a</sup> |
| Analyzed isolates | 3 | 125 | 69 | 98 | 110 | 30 | 30 | 462 |
| pMAB625-1 | 0 | 1 (0.8) | 0 | 0 | 0 | 0 | 0 | 1 (0.2) |
| pMAB625-2 | 0 | 4 (3.2) | 2 (2.9) | 0 | 2 (1.8) | 0 | 2 (6.7) | 10 (2.2) |
| pMAB625-3 | 0 | 16 (12.8) | 5 (7.2) | 8 (8.2) | 14 (12.7) | 9 (30.0) | 1 (3.3) | 53 (11.5) |
| pMAB23 | 1 (33.3) | 0 | 0 | 0 | 1 (0.9) | 0 | 0 | 1 (0.2) |
| pMabS_GZ002 | 1 (33.3) | 0 | 0 | 0 | 1 (0.9) | 0 | 0 | 1 (0.2) |
| pGD69A-2 | 0 | 0 | 0 | 1 (1.0) | 9 (8.2)<sup>b</sup> | 2 (6.7) | 3 (10.0) | 15(3.2) |
| pGD19 | 0 | 0 | 4 (5.8) | 3 (3.1) | 0 | 1 (3.3) | 1 (3.3) | 9(1.9) |
| pJCM30620_1 | 0 | 0 | 1 (1.4) | 0 | 0 | 0 | 1 (3.3) | 2(0.4) |
| 50594_plasmid2 | 0 | 0 | 1 (1.4) | 0 | 0 | 0 | 1 (3.3) | 2(0.4) |
| pGD22-2 | 0 | 0 | 0 | 0 | 6 (5.5) | 4 (13.3)<sup>c</sup> | 1 (3.3) | 11(2.4) |
| pGD42-2 | 0 | 0 | 0 | 1 (1.0) | 12 (10.9)<sup>d</sup> | 4 (13.3)<sup>b</sup> | 1 (3.3) | 18(3.9) |
| pGD25-1 | 0 | 0 | 0 | 0 | 3 (2.7) | 0 | 4 (13.3)<sup>c</sup> | 7(1.5) |
| pGD25-2 | 0 | 0 | 0 | 0 | 1 (0.9) | 0 | 2 (6.7) | 3(0.6) |

Fisher's exact tests were performed to verify regional differences in the frequency of appearance of each plasmid. Corrections for the multiple comparisons were performed using Holm's method.
<sup>a</sup>Total number excludes the number of the type strains.
<sup>b</sup> $p < 0.05$ , compared with Japan
<sup>c</sup> $p < 0.05$ , compared with Japan and Taiwan
<sup>d</sup> $p < 0.05$ , compared with Japan, China and Taiwan

As previous analyses have shown that the pMAB625 plasmids are retained in many clinical isolates, we investigated to what extent other plasmids are retained in the clinical isolates. First, we searched the plasmid sequence in PLSDB as the taxonomic name for “*abscessus”* and got 20 plasmid sequences (Table 3). We removed three plasmids (pGD42-1, pGD69A-1 and FDAARGOS_1640_plasmid) because they were identical to other plasmids with no SV. We selected 20 plasmids shown in Table 3 as “TRUE” for “USE for Reference” together with three pMAB625 plasmids. Then, we mapped the read data of the clinical isolates to these plasmid sequences. Except for the pMAB625 plasmid, three plasmids (pGD69A-2, pGD22-2, and pGD42-2) were detected in more than 10 isolates (Table 4). Seven plasmids (pGD21-1, pGD21-2, pGD22- 1, pGD25-3, pJCM30620_2, BRA100 and 50594_plasmid1) were not detected in our dataset. Statistical analysis showed regional differences in the frequency of appearance of these plasmids (Table 4). pGD69A- 2 was detected more frequently in the US (9/110, 8.2%) than in Japan (0/125, *p*=0.014). pGD22-2 was detected more frequently in Spain (3/40, 13.3%) than in Japan (0/125, *p*=0.018) and Taiwan (0/98, *p*=0.036). pGD42-2 was detected more frequently in Spain (4/30, 13.3%) than in Japan (0/125, *p*=0.017). pGD42-2 was also detected more frequently in the US (12/110, 10.9%) than in Japan (0/125, *p*=0.001), China (0/69, *p*=0.045), and Taiwan (1/98, 1.0%, *p*=0.041). pGD25-1 was detected more frequently in the UK (4/30, 13.3%) than in Japan (0/125, *p*=0.018) and in Taiwan (0/98, *p*=0.036). Moreover, we performed a phylogenetic tree analysis of the chromosomes to determine whether these plasmids are horizontally transferred or whether the same strains harboring the plasmids are spreading (Figure 3). The phylogenetic tree analysis showed that pGD22-2 and pGD42-2 were only detected in the isolates belonging to Clade756. In contrast, pMAB625-3 was detected mainly in the isolates belonging to the four clades (Clade595, 674, 791 and 813). pGD25-1 and pGD69A-2 were detected mainly in the isolates belonging to Clade678 and Clade813. pMAB625-2 and pGD19 were also detected in the isolates belonging to several clades. These results indicated that some plasmids, such as pMAB625-3, were spread across clades and others were restricted to the specific clade.

**Figure 3.**
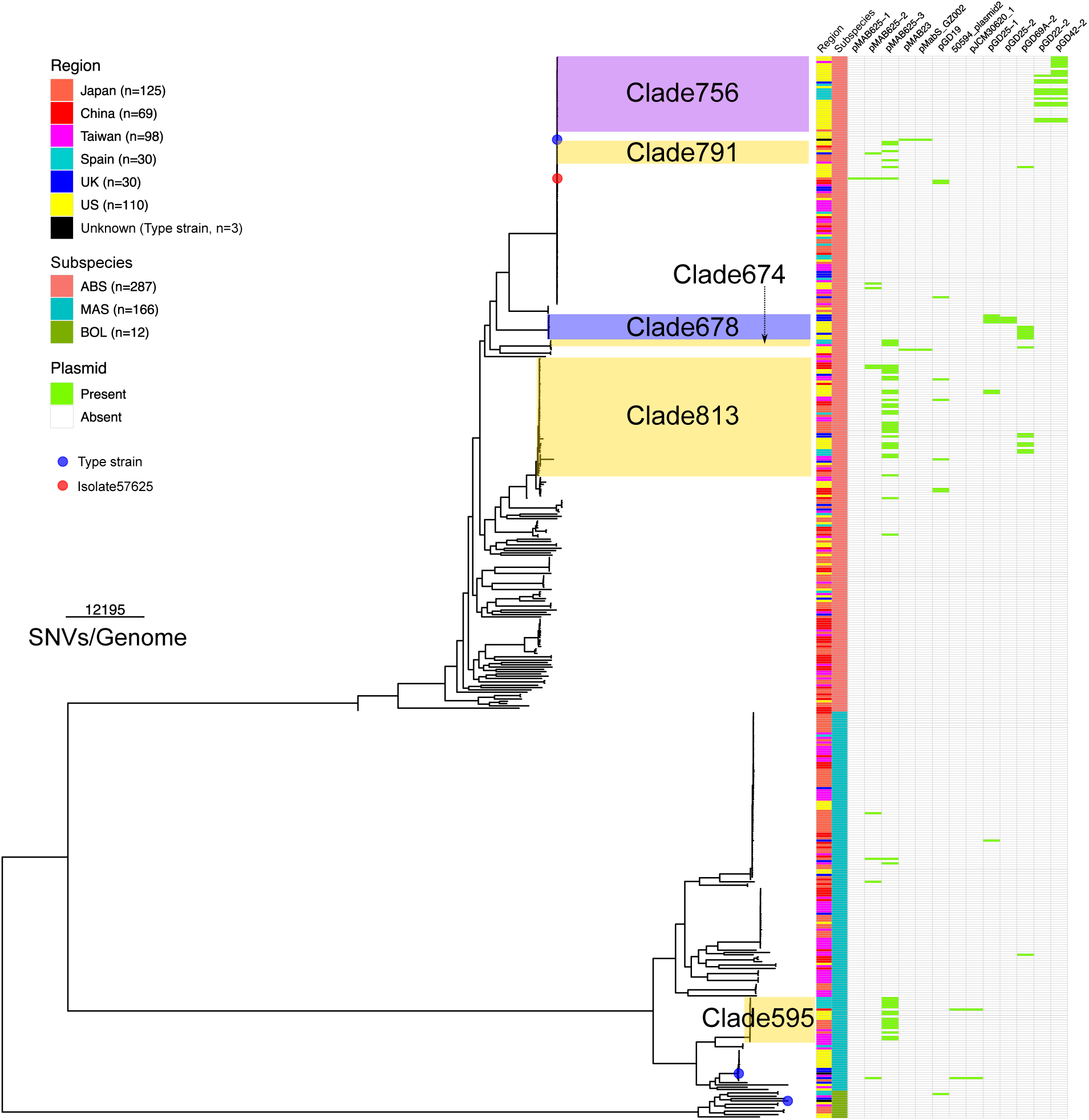
The relation between the phylogenetic distance of host bacterial chromosomes and plasmid distribution in clinical isolates worldwide The phylogenetic tree was created by Gubbins^27^ using chromosome sequences and visualized by ggtree^28^. The read data of the clinical isolates collected worldwide were mapped to pMAB625 and known plasmid sequences to verify the plasmid distribution. Tree-scale represents the number of SNVs per genome.

The phylogenetic tree was created by Gubbins^27^ using chromosome sequences and visualized by ggtree^28^. The read data of the clinical isolates collected worldwide were mapped to pMAB625 and known plasmid sequences to verify the plasmid distribution. Tree-scale represents the number of SNVs per genome.

### Phylogenetic and structural variant analysis of pMAB625 plasmids enabled the strict classification of isolates and prediction of past inter-subspecies contact

So far, we acquired the sequence information for pMAB625-3, which is retained in 53 different clinical isolates, and for pMAB625-2, which is retained in 10 different clinical isolates. Therefore, we examined the genetic relation between the plasmids and investigated whether it could predict host origin and route of transmission. At first, we performed a phylogenetic tree analysis of 53 sequences of pMAB625-3 (Figure 4A). Forty-two sequences (covered by a yellow box in Figure 4A) belonged to the same clade of the sequence of pMAB625-3 in Isolate57625. However, 11 sequences belonged to the different clades. In particular, the pMAB625-3 sequence of SRR7800511 (US, Clade595) was genetically distant from other pMAB625-3 sequences in the isolates belonging to Clace595. The pMAB625-3 sequence of DRR317373 (Japan, Clade791) was also genetically distant from other pMAB625-3 sequences in the isolates belonging to Clade791. These results suggested that plasmid epidemiology, when combined with core-genome analysis, could accurately predict the classification of clinical isolates. In addition, the verification of the mapping results also showed that pMAB625-3 sequences in the six isolates (DRR317373, Japan; SRR18969469, China; SRR7800518, US; SRR7800703, US; SRR7800409, US; SRR7800511, US) were heterogeneous because the unmutated reads and mutated ones were mixed (Figure 4B). Thus, we ran freebays with the parameter “--ploidy 2” to detect SNVs in these mixed reads. Interestingly, the variant hotspot region was located at the CDS of the MMPL family transporter, which was the most reliably highly expressed gene in Isolate57625 and the other four isolates in the RNA-seq analysis. The variant pattern in the MMPL family transporter was classified into three types (Figure 4B). Four sequences (DRR317373, Japan; SRR7800518, US; SRR7800703, US; SRR7800409 US) were classified as Type A and one sequence each was classified as Type B (SRR18969469, China) and Type C (SRR7800511, US), the other 47 sequences had no mutation. There were three missense mutations and 14 or 15 silent mutations in Type A, 20 missense mutations and 43 silent mutations in Type B, and 47 missense mutations and 126 silent mutations in Type C (Table S7). Surprisingly, despite many mutations, no mutation was responsible for the frameshift and nonsense mutations. It was therefore speculated that the mutant MMPL family transporters were being translated into full-length proteins in these isolates. Thus, these data indicated that pMAB625- 3 was spread across strains and regions, each acquiring unique mutations and some clinical isolates retaining pMAB625-3 expressed mutated MMPL family transporters.

**Figure 4.**
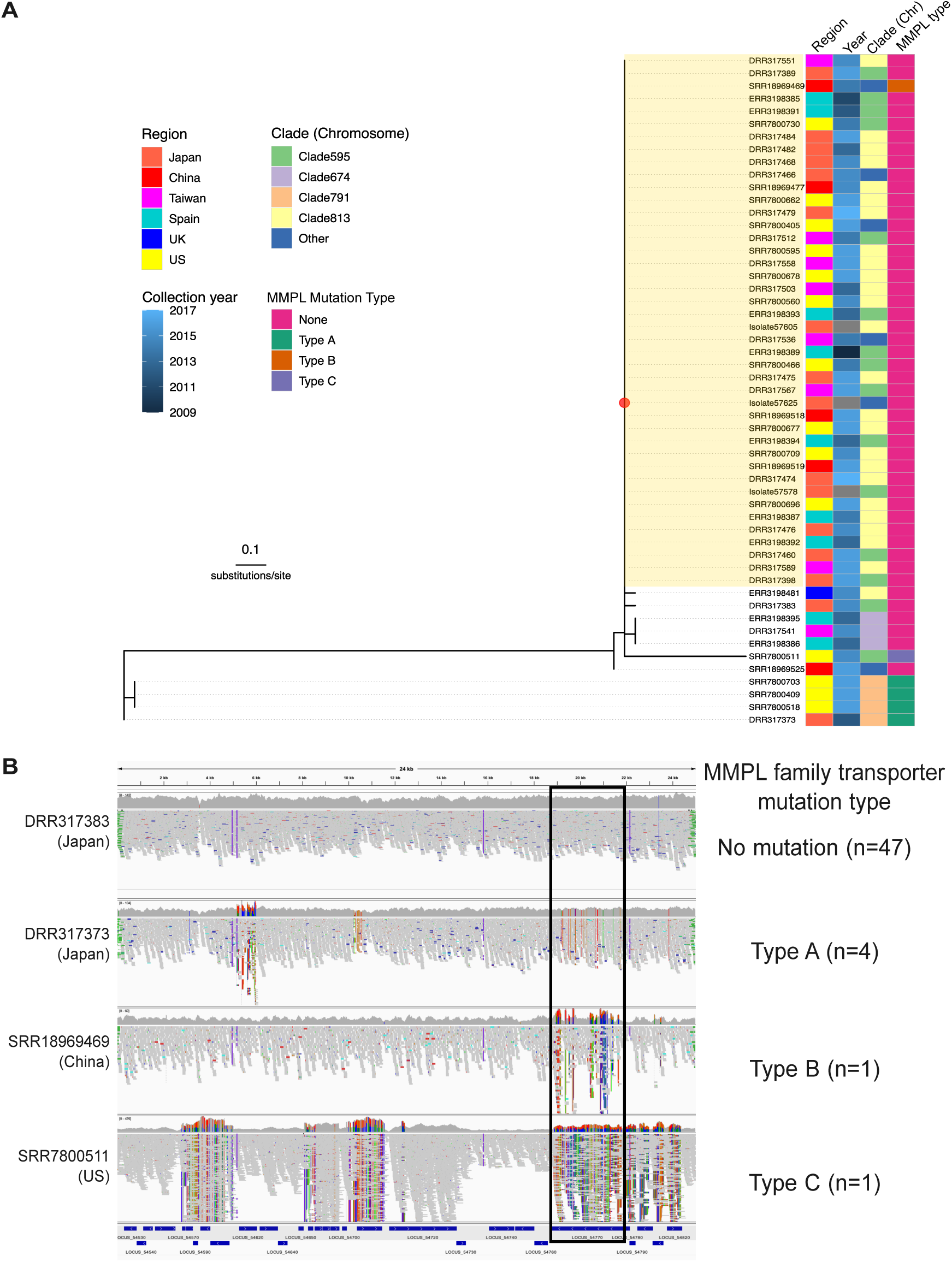
Phylogenetic tree and SNVs analysis of pMAB625-3 (A) Phylogenetic tree of the sequences of pMAB625-3 detected in the 53 isolates. The phylogenetic tree was created using the maximum-likelihood method using iQ-TREE2 based on the core-SNVs data. The sequences belonging to the clade covered by yellow color were identical, although the sequence of SRR18969469 was heterogeneous. The classification of “Clade (Chromosome)” is based on the clade in the phylogenetic tree indicated in Figure 3. pMAB625-3 sequences of the isolates covered by yellow box were identical except for SRR18969469, whose sequence of MMPL family transporter was mixed. (B) The classification of the pMAB625 sequences based on the SNVs. The mapping results of the reads data from each isolate to pMAB625-3 were also visualized by IGV. The mutation hotspot was located at the MMPL family transporter (surrounded by a black line square). The mutation types of the MMPL transporters were classified into three types, and all mutations at this region were missense or silent.

Then, we classified pMAB625-2 sequences in terms of SVs. The sequences of pMAB625-2 were classified into four types (Figure 5A). Assuming that Type I, which was the sequence of pMAB625-2 in Isolate57625, was the base sequence, Type II and III had the large SVs at Region B, and Type II but not Type III had at Region C. In contrast, Type IV had large SVs at Region A and C but not at Region B. Type I, Type III, and Type IV were detected in ABS, and Type II was detected in MAS, except for ERR3198514. In addition, Type III was detected only in the isolates collected in China, and Type IV was detected only in the isolates collected in the US. All pMAB625-2 sequences retained the ESX locus. These results indicated that pMAB625-2 had been horizontally transferred between subspecies and had acquired the SVs unique to each subspecies and region. Next, we analyzed the relation between the phylogenetic distance of hosts’ chromosome sequences and pMAB625-2 type (Figure 5B). The distribution of the pMAB625-2 type was almost consistent with the phylogenetic relation of hosts’ chromosomes. However, the pMAB625-2 type of Isolate57625 and ERR3198415 were different from those of the other isolates belonging to the same clade of phylogenetic tree of hosts’ chromosomes. Interestingly, Type II was detected in four MAS isolates but also in ERR3198415 whose subspecies was ABS. This result indicated the possibility of direct or indirect contact between ERR3198415 and the other four MAS isolates in the past, especially with ERR3198440 isolated from the same region.

**Figure 5.**
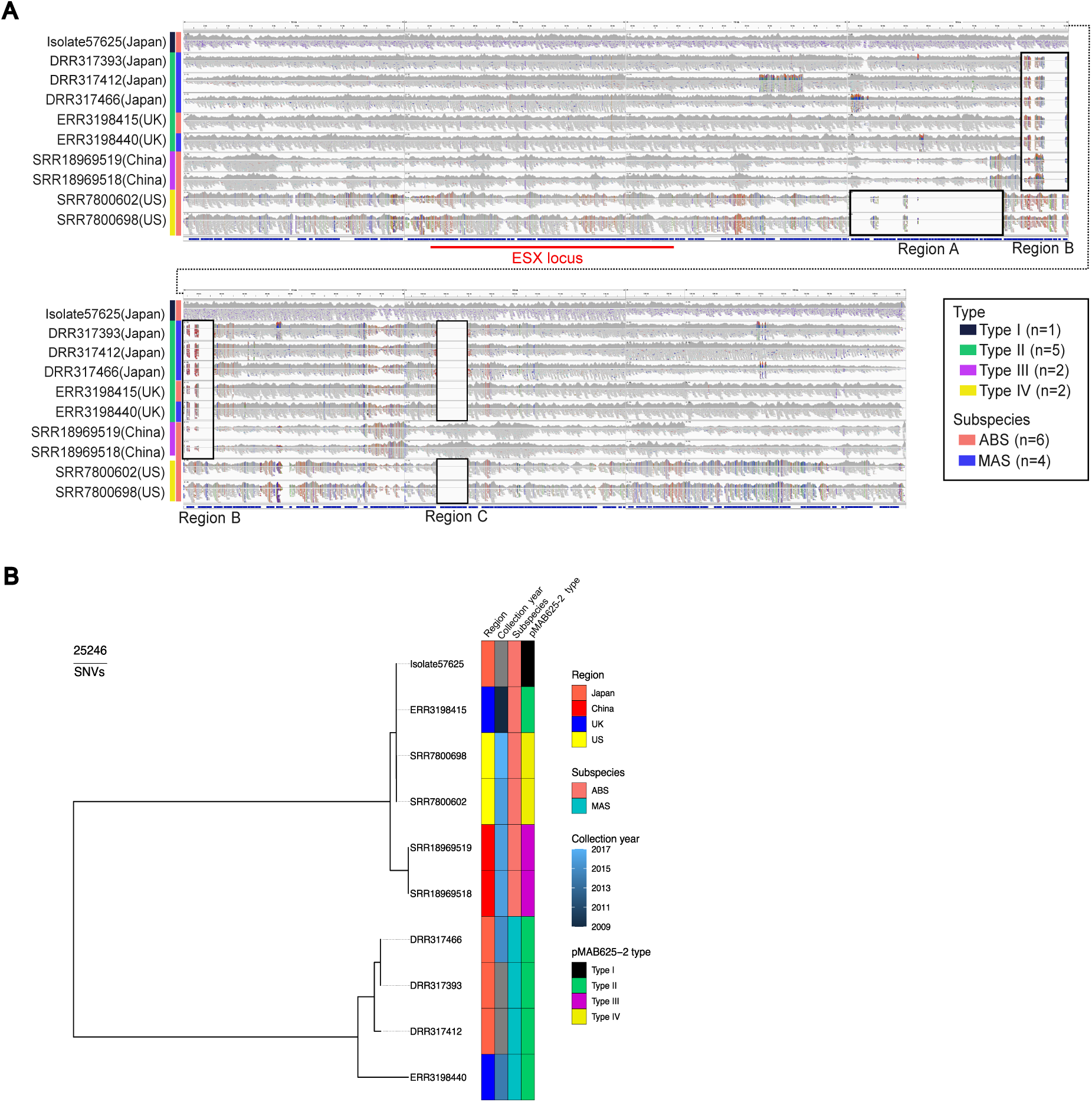
Classification of pMAB625-2 by structural variants and relation with the phylogenetic distance of host bacterial chromosomes (A) The mapping results of the reads data from each isolate to pMAB625-2 were visualized using IGV. Each sequence was classified into 4 types by the pattern of structural variants (surrounded by black line squares). (B) Relation between the phylogenetic distance and pMAB625-2 distribution.

Finally, we investigated the distribution in our dataset regarding the plasmids identified in other NTMs except for *M. abscessus* to verify whether plasmids transferred between *M. abscessus* and other NTM species. Recently, Wetzstein *et al*. reported clinical and genomic features in Europe about *Mycobacterium avium* complex (MAC)^29^, the most frequently detected NTM globally^30,31^. In the report, they investigated the distribution of the known 152 plasmids registered in PLSDB and showed that plasmids identified in *M. abscessus*, including pGD69B-1 (identical to pGD69A-1) and pGD42-1, which were identical to pMAB625- 3, were not detected in their dataset. Therefore, we investigated the distribution of these 152 plasmids in our dataset using our methods in this study. As a result, we detected the same plasmids indicated in Table 3 but did not other plasmids identified in other NTM except for pMN23 (accession number: NC_010604.1) and pMFLV03 (accession number: NC_009341.1) in two and one isolates, respectively (Table S8). pMN23 was almost identical to pMAB23 (ANI: 99.9%), so pMN23 was detected redundantly in ATCC 19977 and SRR7800408, which had pMAB23. Interestingly, pMFLV03 was detected in ERR3198408 (covered genome region: 78.7%, depth of coverage: 73.1x). pMN23 had originally identified in *Mycobacterium marinum*^32^ and pMFLV03 had identified in *M. gilvum* PYR-GCK (according to the GenBank registry information), indicating that these plasmids could be present across NTM species. These results suggested that plasmid distribution is strictly different from *M. abscessus* and other NTMs except for very few cases.

## DISCUSSION

Plasmids are mobile genetic elements that can move between different bacterial strains and alter host characteristics, such as virulence, and antimicrobial susceptibility. Plasmids are horizontally transferable among bacteria by transformation and conjugation. Thus, plasmid epidemiology, in conjunction with core-genome analysis, may help to infer past contact between bacteria. In this study, we showed that the multidrug-resistant clinical isolates of ABS isolated from a patient with osteomyelitis had three plasmids (pMAB625-1, pMAB625-2 and pMAB625-3) that the type strain did not harbor. Additionally, analysis of the clinical isolates of *M. abscessus* identified worldwide indicated that pMAB625 plasmids were horizontally transferred between regions and subspecies and that it is sometimes possible to predict past contact between bacteria by tracing the phylogenetic relation of plasmids. Dedrick *et al*. reported that pGD69-1 (pGD69A-1), identical to pMAB625-3, is the same as the other five plasmids^33^. Lewin *et al*. identified pMabs-09-13 through WGS of the clinical isolates of *M. abscessus* within cystic fibrosis patients and reported that pMabs-09-13 exhibits an identity of 99.86% to pGD42-1, pGD69A-1, and pGD69B-1^34^. These reports suggested that pMAB625-3 is spreading among many other isolates of *M. abscessus*, but the detailed distribution of pMAB625 plasmids in the world and other subspecies has remained unclear. Our study shows that pMAB625-3 is the plasmid spreading beyond regions and subspecies in the largest number of clinical isolates of *M. abscessus*. This allowed us to investigate the possibility of speculating the precise classification of bacteria and the past contact between them. The phylogenetic tree analysis of pMAB625-3 based on the SNVs of plasmids (Figure 4A) indicated that the SNV pattern was correlated with the phylogenetic relation of the hosts’ chromosome sequences; however, a unique variant (SRR7800511, US) different from the other sequences in the clade was found. This indicates that it is possible to predict the bacterial contact using the distribution of the unique variant like SRR7800511. In the case of pMAB625-2, it is actually indicated that the past contact between the different subspecies is predictable based on the relation between the phylogenetic tree of the hosts’ chromosomes and the distribution of pMAB625-2 type.

As important findings, we found that 2.2% (10/462) and 11.5% (53/462) of the clinical isolates worldwide had pMAB625-2, and pMAB625-3, respectively (Table 4). This result provides an epidemiological insight into the spread of plasmids in *M. abscessus* beyond regions and strains. Other than person-to-person transmission of *M.abscessus*, this global spread of pMAB625 plasmids may be mainly due to increased global logistical activity and human mobility with associated interference of human and microbial habitats^35^. The habitat of *M. abscessus* is soil and water systems, and it is also possible that increased flooding due to recent climate change may have increased the chance of contact between the different strains. Analysis of plasmid distribution revealed that some plasmids (pMAB625-2, pMAB625-3, pGD19, pGD25-1 and pGD69A-2) have been widely spread between strains, regions and subspecies, while others (pGD22-2 and pGD42-2) have been restricted to specific strains and regions (Figure 3). Therefore, it is speculated that regarding pGD22-2 and pGD42-2, the identical isolates retaining these plasmids had been spread within or between the regions, whereas, regarding pMAB625-2, pMAB625-3, pGD19, pGD25-1 and p69A-2, plasmids themselves had been spread between different strains, subspecies and region through horizontal transfer. It is unknown what factors determine these differences in plasmid distribution. It is possible that the factors encoded on the plasmid or the periods from the emergence were different among the plasmids. In addition, we also showed the plasmid distribution was different from *M. abscessus* and MAC (Table S8), indicating that incompatibility also exists in the plasmids present in NTM, strictly distinguishing *M. abscessus* from other NTMs except for very few cases, such as pMN23 and pMFLV03.

We identified various factors associated with bacterial virulence and multidrug resistance on the pMAB625 plasmids. The components of the ESX secretion system, the MMPL family transporter, and the components of the TA system were encoded on the plasmids, and these genes were highly expressed in the clinical isolates compared to the type strain. The ESX secretion system is involved in the export of the various pathogenic proteins and metal ions associated with proliferation, survival in the cytosol of macrophage, and conjugation. It has also been reported that ESX-1 components are required for sliding motility and biofilm formation in *M. avium*^20^. Therefore, given the close genetic distance between ESX-1 and ESX-P cluster 5, which include the components of the ESX secretion system on the pMAB625-1 and pMAB625-2 (Figure 1B), the components of ESX-P cluster 5 may be involved in biofilm formation. In addition, MMPL family transporters can transport antibiotics across the cell membrane. It has been reported that the increased expression of MMPL5 is associated with decreased susceptibility to bedaquiline, and clofazimine in *M. tuberculosis*^36,37^, bedaquiline, and clofazimine in *M. intracellulare*^38^, and thiactazone derivative in *M. abscessus*^39^. The function of the MMPL family transporter encoded on the pMAB625-3 is unknown; however, the presence of mutational hotspots (Figure 4B) indicates that this MMPL transporter may be compatible with a variety of substrates, including antimicrobial agents. Therefore, our results suggest that Isolate57625 enhanced defence mechanisms against antibiotics through the acquisition of the ESX secretion system and transporters, accompanied by changes in biofilm barriers, and transport of antibiotics. These results suggest that pMAB625 plasmids appeared to be a cause of the low susceptibility to carbapenem antibiotics in the clinical isolates compared with the type strain; however, further studies will be essential to determine whether these plasmids are the main cause of AMR.

In conclusion, we reported the distribution of plasmids in *M. abscessus* and found that pMAB625-3 is the most widely distributed plasmid in the clinical isolates of *M. abscessus* worldwide. In addition, phylogenetic tree analysis and mutational analysis of the pMAB625 plasmids partially predict past bacterial contact. These findings provide a new perspective on the acquisition of genomic diversity, the origin and transmission route of *M. abscessus.* These findings will also lead to the development of effective countermeasures to identify pathways of bacterial transmission and prevent serious bacterial outbreaks.

### Limitations of the study

Although it is assumed that phylogenetically close plasmids with similar SNV and SV patterns were spread via bacterial contact, it cannot be excluded that they may have acquired identical mutation patterns in their respective clinical isolates without bacterial contact. To clarify this, it is necessary to calculate the mutation rate of the plasmid and compare the mutation acquisition period with the interval of appearance in the different strains. In addition, future studies are required to verify whether the pMAB625 plasmids induce multidrug resistance in *M. abscessus* by plasmid transfection.

## RESOURCE AVAILABILITY

### Lead contact

Further information and requests for resources and reagents should be directed to and will be fulfilled by the lead contact, Masashi Toyoda.

### Materials availability

This study did not generate new unique reagents.

### Data and code availability

- All raw sequence data generated in this study have been deposited in the DDBJ database at https://www.ddbj.nig.ac.jp/ under the BioProject accession number PRJDB16220. The complete genome sequences of the isolates are available in the DDBJ database. They can be accessed using the accession numbers, AP028613, AP028614, AP028615, and AP028616 for Isolate57625; AP028617, AP028618, AP028619, and AP028620 for Isolate57629; AP028621, AP028622, AP028623, and AP028624 for Isolate57630; AP028625, AP028626, AP028627, and AP028628 for Isolate57596; AP028629, AP028630, AP028631, and AP028632 for Isolate57626, for the chromosome and three plasmids in each isolate.
- The analyzed data of RNA-seq have been deposited in the Genomic Expression Archive at https://www.ddbj.nig.ac.jp/gea/ under the accession number E-GEAD-827.
- All data reported in this paper will be shared by the lead contact upon request.
- This paper does not report original code.
- Any additional information required to reanalyze the data reported in this paper is available from the lead contact upon request.

## ACKNOWLEDGMENTS

The authors thank the staff of the microbiology laboratory of Dokkyo Medical University Hospital for performing identification and drug susceptibility tests, and Dr. Yoshishige Masuda and Dr. Takashi Inamatsu at TMIG for supervising in clinical practice. Computations were partially performed on the NIG supercomputer at ROIS National Institute of Genetics. This work was supported by Japan Society for the Promotion of Science (JSPS) KAKENHI Grant Number JP19K08938.

## AUTHOR CONTRIBUTIONS

Conceptualization,K.O., A.Y.; Formal analysis, K.O.; Methodology, K.O., A.Y., and Hiroshi Koganemaru; Data curation, K.O.; Investigation,K.O., Keisuke Kamada, and Ken Kikutchi; Validation, K.O.; Writing-original draft, K.O. and A.Y.; Writing-review & editing, K.O., Y.A. and M.T.; Funding acquisition, A.Y. and Ken Kikuchi.; Patient care and resources, Hironobu Kitazawa, Y.I., T.S. and K.W.; Supervision, M.T.

## SUPPLEMENTAL INFORMATION

**Document S1. Figures S1–S6**

**Table S1. Primer sequences used in this study**

**Table S2. Genbank accession numbers of the genome and plasmid sequences in this study**

**Table S3. Gene lists on chromosome which were acquired, lost and mutated in Isolate57625 compared with ATCC19977**

**Table S4. Gene lists which were acquired, lost and mutated in Isolate57625 compared with ATCC19977**

**Table S5. Homology search results of genes on pMAB625 plasmids against genes of Mycobacterium**

**Table S6. The results of RNA-seq analysis**

**Table S7. The mutation type of MMPL family transporter**

**Table S8. The analysis of distribution regarding NTM plasmids in the clinical isolates of *M. abscessus***

---

**STAR★METHODS**

## KEY RESOURCES TABLE

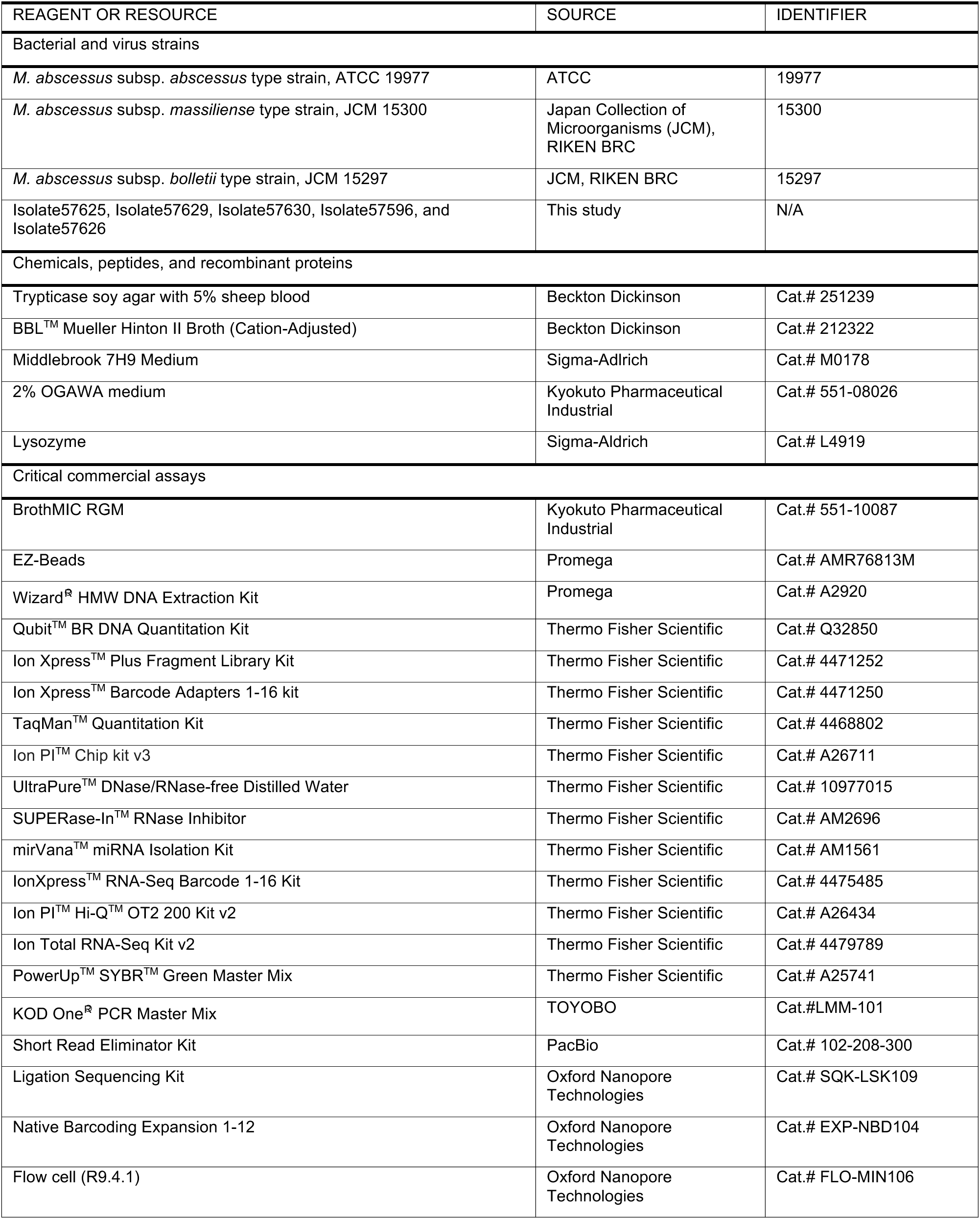
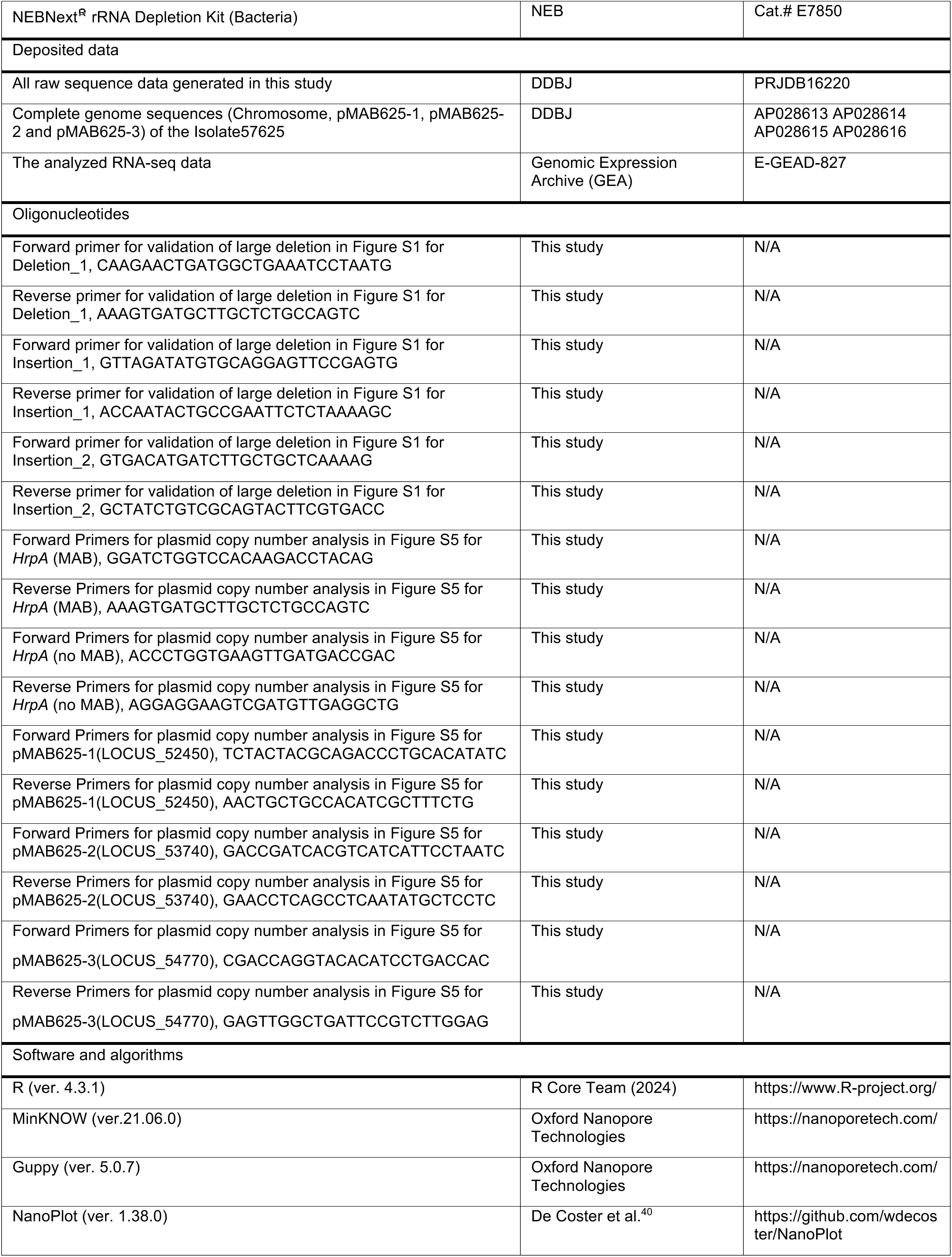
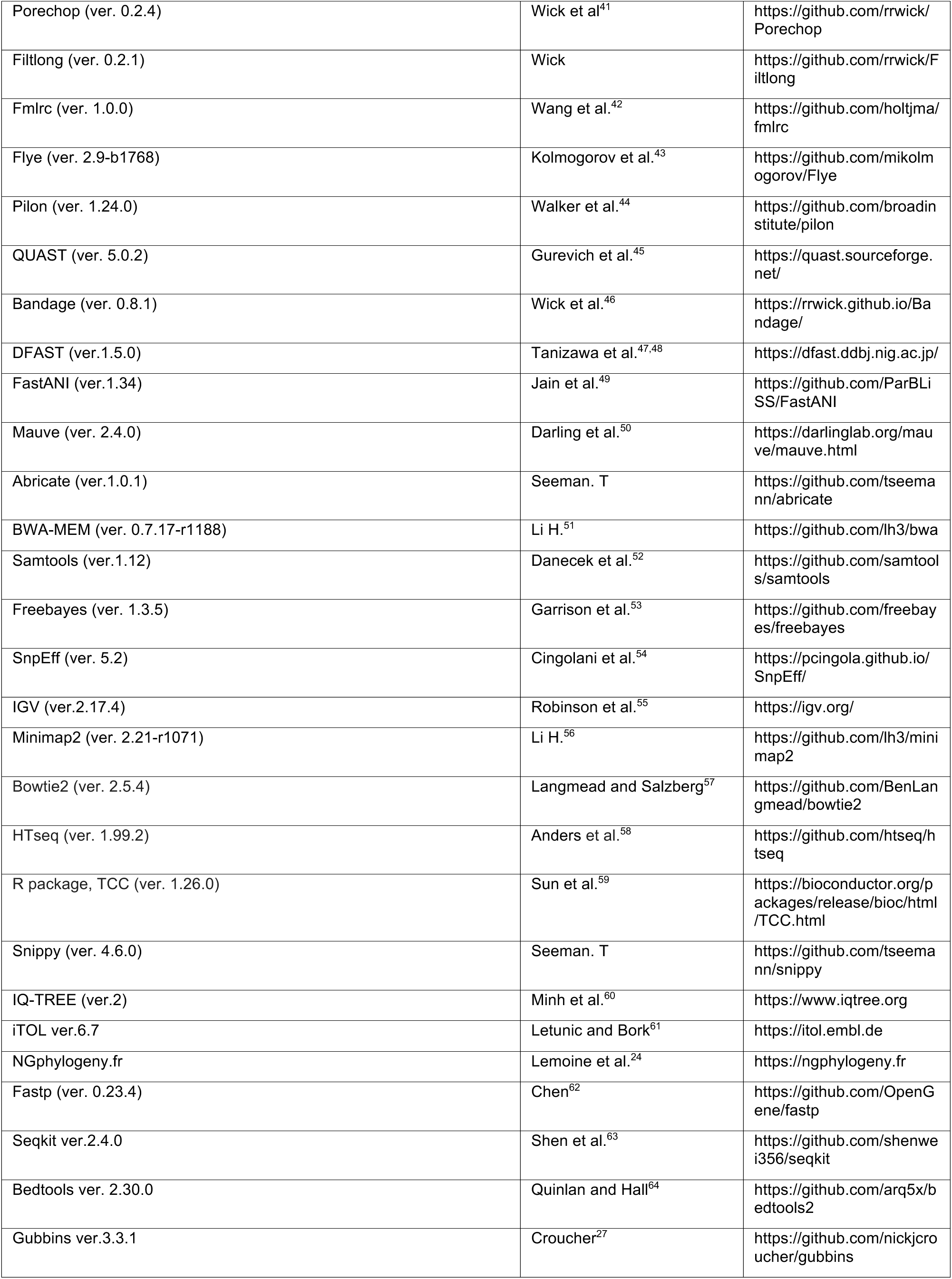
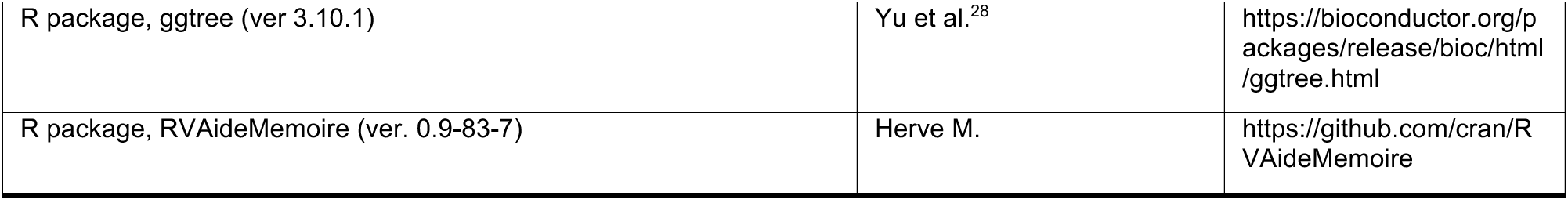

## EXPERIMENTAL MODEL AND STUDY PARTICIPANT DETAILS

### Clinical isolates and type strain

Ethical approval for this study was obtained from the Ethics Committee of Dokkyo Medical University (approval number 28045). Regarding Isolate57625, Isolate57629, Isolate57630, Isolate57596, and Isolate57626, specimens were obtained from a patient with osteomyelitis at the five time points in six months indicated in Table 1. Each strain was isolated from the specimen at Shizuoka Children’s Hospital for diagnosis. The other clinical isolates used for PCR or WGS were stored at Tokyo Metropolitan Institute for Geriatrics and Gerontology or Tokyo Women’s Medical University or Dokkyo Medical University^25^. The type strain of ABS, ATCC 19977, was purchased from ATCC (Manassas, VA, US). The type strain of MAS, JCM 15300, and the type strain of BOL, JCM 15297, were provided by Japan Collection of Microorganisms, RIKEN BRC which is participating in the National BioResource Project of the MEXT, Japan.

## METHOD DETAILS

### Growth curve

The type strain and the five clinical isolates were inoculated onto trypticase soy agar with 5% sheep blood (Beckton Dickinson, Franklin Lakes, NJ, US) and incubated at 30 °C for five days. Next, some colonies were inoculated into BBL^TM^ Mueller Hinton II Broth (Cation-Adjusted) (Beckton Dickinson) and incubated overnight at 30°C. These culture media were measured for absorbance (OD_600_), from which they were prepared to 0.1 by medium. Seventy microliter of the adjusted culture solution was inoculated into 7 mL of BBL^TM^ Mueller Hinton II Broth (Cation-Adjusted) and incubated at 30 °C for eight days with agitation (200 rpm). OD_600_ was measured on days one to eight after inoculation. Statistical analysis was performed using R ver. 4.3.1.

### Antimicrobial susceptibility testing

The type strain and the clinical isolates were subcultured on 2% OGAWA medium (Kyokuto Pharmaceutical Industrial, Tokyo, Japan) at 30 °C for five days. Antimicrobial susceptibility testing was performed using BrothMIC RGM (Kyokuto Pharmaceutical Industrial) following CLSI M24 3rd ed^65^. MICs were determined as the minimum concentration at which bacteria had not grown on day four (all antibiotics) and day 14 (clarithromycin). The test was repeated three times and all MICs were confirmed to be the same between the experiments, except for imipenem (IPM) for the isolates. For IPM, the MICs of the type strain were confirmed to be the same between all three experiments, while for the MICs of Isolate57625 and Isolate57626, they were confirmed to be the same between two out of three experiments, as indicated in Table 1.

### Mycobacterial DNA isolation and whole-genome sequencing

The type strain and the clinical isolates of *M. abscessus* were stored in frozen stock containing skim milk and 5% (v/v) glycerol. Each strain was inoculated from frozen stocks into Middlebrook 7H9 Medium (Sigma-Aldrich, St. Louis, MO, US) containing 10% (v/v) oleic acid-albumin-dextrose-catalase and 0.2% (v/v) glycerol and cultured at 30 °C for six days with agitation (200 rpm). Bacterial genomic DNA for short-read sequencing was extracted as previously reported^66^. The quality of the extracted DNA was confirmed by agarose gel electrophoresis, and the concentration was measured using Qubit^TM^ BR DNA Quantitation Kit (Thermo Fisher Scientific, Waltham, MA, US). Genomic DNA was fragmented, and the sequencing library was prepared using the Ion Xpress^TM^ Plus Fragment Library Kit and the Ion Xpress^TM^ Barcode Adapters 1-16 kit (Thermo Fisher Scientific) according to the manufacturer’s instructions. The quantity of sequencing library was assessed using the Ion Library TaqMan^TM^ Quantitation Kit (Thermo Fisher Scientific). Emulsion PCR for sequence template synthesis was performed with the Ion PI^TM^ Hi-Q^TM^ OT2 200 Kit v2 using the Ion One Touch^TM^ II System and the Ion One Touch^TM^ II ES (Thermo Fisher Scientific). Sequencing was performed using the Ion PI^TM^ Chip kit v3 and the Ion Proton^TM^ System (Thermo Fisher Scientific). Data were collected using the Torrent Suite v5.0.5 software (Thermo Fisher Scientific).

High-molecular-weight genomic DNA for long-read sequencing was extracted as follows. The bacterial pellet was collected by centrifugation and homogenized using EZ-beads (Promega, Madison, WI, US) and Shakeman 6 (Bio Medical Science, Tokyo, Japan). The homogenized pellet was lysed with Lysozyme (Sigma-Aldrich) at 37 °C for one hour. Next, DNA was extracted using Wizard^○R^ HMW DNA Extraction Kit (Promega) following the manufacturer’s instructions. Fragmented DNA in the extracted DNA was removed using the Short Read Eliminator Kit (PacBio, Menlo Park, CA, US). The sequencing library for long-read sequencing was prepared using the Ligation Sequencing Kit (SQK-LSK109, Oxford Nanopore Technologies, Oxford, UK) and the Native Barcoding Expansion 1-12 (EXP-NBD104, Oxford Nanopore Technologies). In brief, the barcode sequences for multiplex sequencing were attached to the ends of the HMW genomic DNA. In addition, the sequencing adapters were ligated to the end of HMW genomic DNA. The prepared DNA was loaded into a flow cell (FLO-MIN106, Oxford Nanopore Technologies) attached to MinION Mk1B (Oxford Nanopore Technologies). Sequencing data were acquired using MinKNOW ver.21.06.0 (Oxford Nanopore Technologies), and base calling was performed using Guppy ver. 5.0.7 (Oxford Nanopore Technologies) with high accuracy mode using the National Institute of Genomics supercomputer system (NIG, Shizuoka, Japan). Data quality was confirmed using NanoPlot ver. 1.38.0^40^ and adapter sequences were trimmed using Porechop ver. 0.2.4^41^. Raw reads shorter than 1000 bp (Isolate57625, Isolate57629, Isolate57630, Isolate57596) or 5000 bp (Isolate57626) were removed using Filtlong ver. 0.2.1 (https://github.com/rrwick/Filtlong). Filtered reads were corrected using Fmlrc ver. 1.0.0^42^ with the data of short-read sequencing acquired by the Ion Proton^TM^ Sequencer. *De novo* assembly was performed using Flye ver. 2.9-b1768^43^ with corrected long-reads. Assembled genome sequences were polished using Pilon ver. 1.24.0^44^, evaluated using QUAST ver. 5.0.2^45^, and visualized using Bandage ver. 0.8.1^46^. Genome annotation was performed using DFAST ver.1.5.0^47,48^.

### Plasmid sequence analysis

Known plasmid sequences were downloaded from PLSDB^14^ by the taxonomic search for “*abscessus*”. Average nucleotide identity (ANI) was calculated using FastANI ver.1.34^49^ and structural variants (SVs) were analyzed using Mauve ver. 2.4.0^50^.

Homology search was performed by blast+^67^ using the amino acid sequence as the query or by Abricate ver.1.0.1 (http://github.com/tseemann/abricate) using the nucleotide sequences as the query. NCBI AMRFinderPlus^68^, CARD^69^, Resfinder^70^, ARG-ANNOT^71^, VFDB^72^, PlasmidFinder^73^, EcOH^74^, and MEGARes 2.00^75^ were used as the database for the search of AMR genes. Mycobrowser^17^ was used as the database for the homology search of the mycobacterium genes.

### SNVs and SVs analysis

Short-read data acquired by the Ion Proton^TM^ Sequencer were mapped using BWA-MEM ver. 0.7.17-r1188^51^ against the reference sequence (CU458896.1, CU458745.1). Output BAM files were sorted using samtools ver.1.12^52^. Valiant calling was performed using Freebayes ver. 1.3.5^53^, and variants were annotated using SnpEff ver. 5.2^54^ with annotated genome sequence. The positions of single nucleotide variants (SNVs) were visualized using IGV^55^ (Broad Institute, Cambridge, MA, US). Assembled genome sequences were aligned using Mauve ver. 2.4.0^50^ for SVs analysis. Mapping the short-and long-read to the assembled genome sequence using minimap2 ver. 2.21-r1071^56^ was performed for the validation of assembly.

PCR for the validation of SVs was performed using KOD One^○R^ PCR Master Mix (TOYOBO, Osaka, Japan) and the primers indicated in Table S1. Amplified DNA was confirmed using agarose gel electrophoresis following staining with SYBR^TM^ Safe Gel Stain (Thermo Fisher Scientific).

### RNA-seq analysis

Total RNA was extracted as follows: The bacterial pellet was collected by centrifugation and suspended in UltraPure^TM^ DNase/RNase-free Distilled Water (Thermo Fisher Scientific) and homogenized using EZ-beads (Promega) and Shakeman 6 (Bio Medical Science). SUPERase-In^TM^ RNase Inhibitor (Thermo Fisher Scientific) was added to the homogenized pellet. Next, the homogenized pellet was lysed by Lysozyme (Sigma-Aldrich) at 37 °C for an hour. Total RNA was purified by mirVana^TM^ miRNA Isolation Kit (Thermo Fisher Scientific). The size and quality of total RNA were assessed using the Agilent 2100 Bioanalyzer (Agilent Technologies, Santa Clara, CA, US) and the RNA 6000 Nano Kit (Agilent Technologies). rRNA was depleted using the NEBNext^○R^ rRNA Depletion Kit (Bacteria) (NEB, Ipswich, MA, US) from 500 ng of total RNA. The whole transcriptome library was prepared using the Ion Total RNA-Seq Kit v2 (Thermo Fisher Scientific) as follows: First, the rRNA-depleted RNA was fragmented by RNase III at 37 ℃ for 10 minutes and purified using the Magnetic Beads Cleanup Module. Next, adapter sequences were ligated and reverse transcription was performed. The whole transcriptome library was amplified and barcoded using the IonXpress^TM^ RNA-Seq Barcode 1-16 Kit (Thermo Fisher Scientific). The quantity of the amplified whole transcriptome library was assessed using the Ion Library TaqMan^TM^ Quantitation Kit (Thermo Fisher Scientific). Emulsion PCR for sequence template synthesis was performed using the Ion PI^TM^ Hi-Q^TM^ OT2 200 Kit v2 with the Ion One Touch^TM^ II System and the Ion One Touch^TM^ II ES (Thermo Fisher Scientific). Sequencing was performed using the Ion Proton^TM^ System with Ion PI^TM^ Chip kit v3 (Thermo Fisher Scientific). Data was collected using the Torrent Suite ver. 5.0.5 software (Thermo Fisher Scientific). Sequence reads were mapped using Bowtie2 ver. 2.5.4^57^ against the assembled genome sequence of Isolate57625. Mapped reads were counted using HTseq ver. 1.99.2^58^. Merged transcript count data were normalized using TCC ver. 1.26.0^76^ and differentially expressed genes (DEG) were detected under false discovery rate (FDR) < 0.01.

### Phylogenetic tree analysis

The genome sequences of the various clinical isolates shown in Figure S2C for phylogenetic tree analysis were obtained from the NCBI database and are listed in Table S2. Core-genome SNVs were detected using Snippy ver. 4.6.0 (https://github.com/tseemann/snippy) with each assembled genome sequence and the type strain (ATCC 19977, CU458896.1) as the reference. The concatenated core-genome SNVs were also aligned using the snippy-core script. The phylogenetic tree was constructed using IQ-TREE ver.2^60^ using the maximum likelihood criterion and rendered using iTOL ver.6.7^61^. Phylogenetic tree analysis of pMAB625 sequences was performed by the same method. For the phylogenetic tree analysis regarding the components of the ESX secretion system, the amino acid sequences of ESX loci (*eccB*, *eccC*, *eccD*, *mycP*, and *eccE*) were concatenated and used as input for NGphylogeny.fr^24^ using the PhyML+SMS inference method.

### PCR for plasmid detection and copy number analysis

PCR was performed using KOD One^○R^ PCR Master Mix (TOYOBO) and the primers indicated in Table S1. Amplified DNA was confirmed using agarose gel electrophoresis. The plasmid copy number was analyzed by qPCR using PowerUp^TM^ SYBR^TM^ Green Master Mix (Thermo Fisher Scientific). qPCR was performed using the QuantStudio^TM^ 5 Real-Time PCR System (Thermo Fisher Scientific). Five nanogram of genomic DNA was used for a 20 μL reaction volume of qPCR. The plasmid copy number is shown as a relative value normalized using the copy number of *HrpA*, which is present in one copy on the chromosome.

### Mapping the read data of various clinical isolates to the genome sequence of Isolate57625

Raw read data of *M. abscessus* clinical isolate in Japan, Taiwan^7^, China^26^, the US, the United Kingdom (UK) and Spain were obtained from the NCBI sequence reads archive (SRA). Raw read data were trimmed using fastp ver. 0.23.4^62^ and reduced using seqkit ver.2.4.0^63^ to 200 Mb for the Ion Proton^TM^ single-end data and 100 Mb for the Illumina pair-end data to equalize the depth of coverage among isolates. Reads were mapped using BWA-MEM ver. 0.7.17-r1188 against the chromosome and plasmid sequences of Isolate57625 and other plasmids which were hit using PLSDB by the taxonomic searching of “*abscessus*” with removing the redundant plasmid sequences (ANI (%)>97, without large SV (>8 kbp)). The depth of coverage was calculated using Qualimap ver. 2.2.2-dev^77^. The covered genome region was calculated using bedtools ver. 2.30.0^64^. We considered the isolate whose depth of coverage was higher than 10x and the covered genome region was higher than 75% as the plasmid-harboring isolate. The 152 sequences for the analysis of the distribution of plasmids identified in other NTM and previously reported^29^ were listed in Table S2. For the phylogenetic tree analysis, the aligned assumed chromosome sequences using Snippy-core script were used as input for Gubbins ver.3.3.1^27^. The phylogenetic tree was rendered using ggtree ver 3.10.1^28^.

## QUANTIFICATION AND STATISTICAL ANALYSIS

Statistical analysis in this study were performed using R ver. 4.3.1. Statistical significance in Figure S1 were calculated using one-way ANOVA followed by Dunnett’s test and data were represents the mean of the three independent experiments and the error bar represents the standard deviation. Statistical analysis of the number of clinical isolates retaining plasmid in Table 4 was performed using Fisher’s exact test with Holm’s method for correction of multiple comparisons using RVAideMemoire ver. 0.9-83-7 (https://github.com/cran/RVAideMemoire).

