## Supplementary material for "Genetic analysis of *Mycobacterium abscessus* reveals genomic diversity linked to global plasmid distribution": Document S1

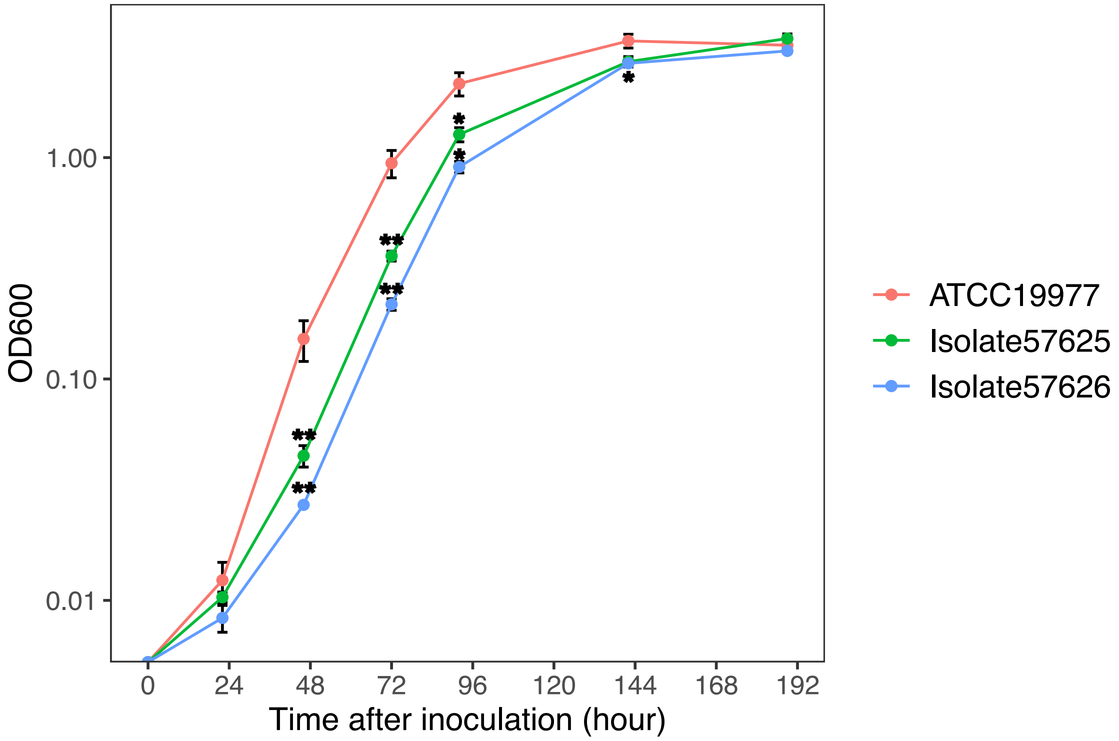


**Figure S1. Growth curve of the type strain and the isolates**

The type strain (ATCC 19977) and the isolates were inoculated with the same seeding concentration and cultivated at 30 ° C. $OD_{600}$ was measured at each time point. Data represents the mean of the three independent experiments and the error bar represents the standard deviation. Asterisks represent the statistical significance between ATCC 19977 and each isolate. *p*-value were calculated using one-way ANOVA followed by Dunnett's test (**p*<0.01, ***p*<0.001).

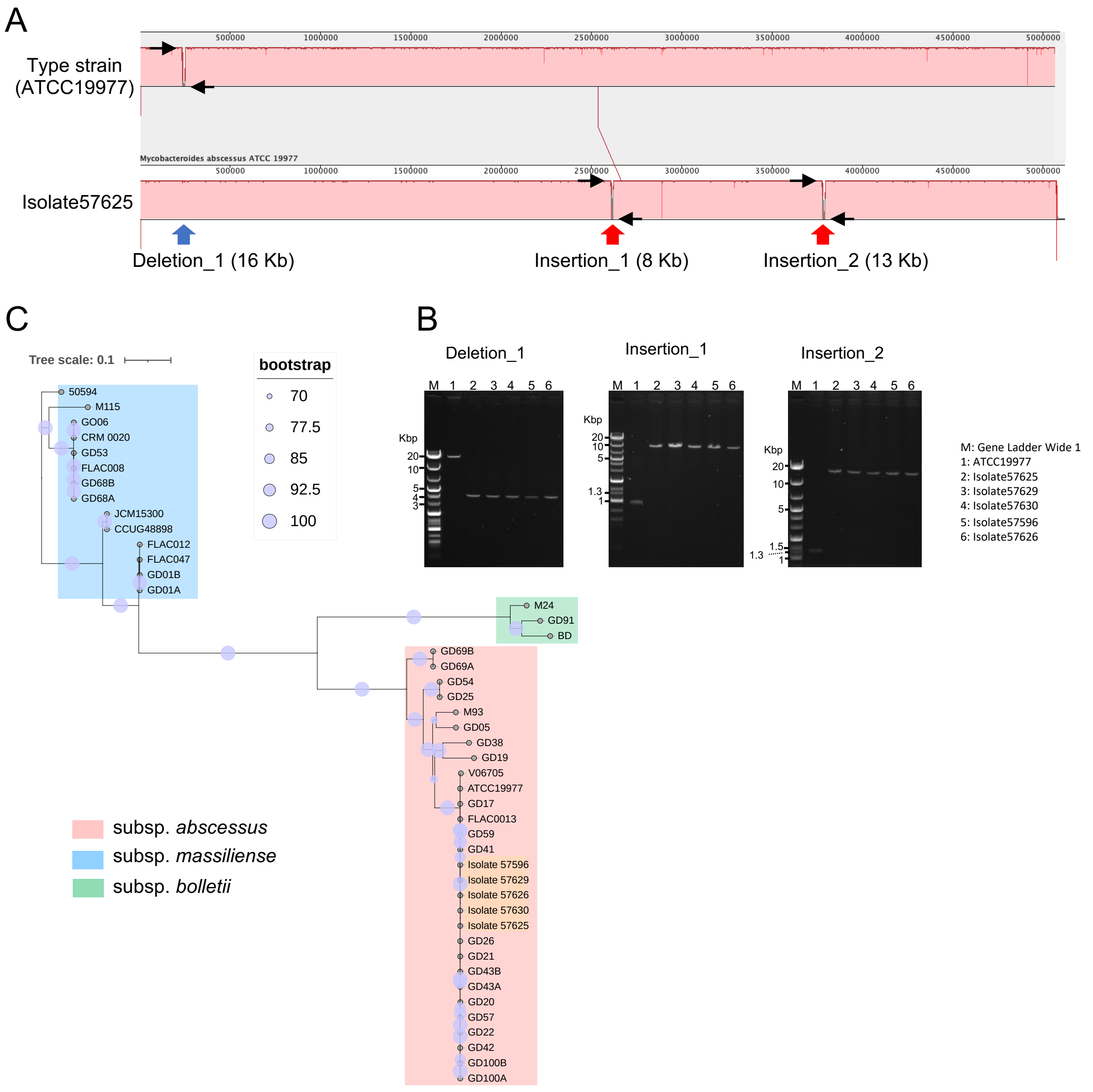


**Figure S2. Comparison of the chromosome sequence of Isolate57625 with the type strain and other isolates**

(A) The alignment of chromosome sequence of the type strain (ATCC 19977) and Isolate57625 detected three large structural variants (SVs). One deletion and two insertions were detected in the chromosome of Isolate57625 compared to that of ATCC 19977. Black arrows indicated the primer region used for the validation of the SVs.

(B) The validation of SVs using PCR. PCR was performed using the genomic DNA and the primers flanking the SVs.

(C) The phylogenetic tree of the relevant isolates and other clinical isolates was constructed using the core single nucleotide variants (SNVs) detected in each chromosome sequence by the maximum likelihood method. Zero point one of the tree scale represents 10 % differences in the core SNVs. The bootstrap value (%) represents the probability that the same branch was observed when a phylogenetic tree was generated 1000 times.

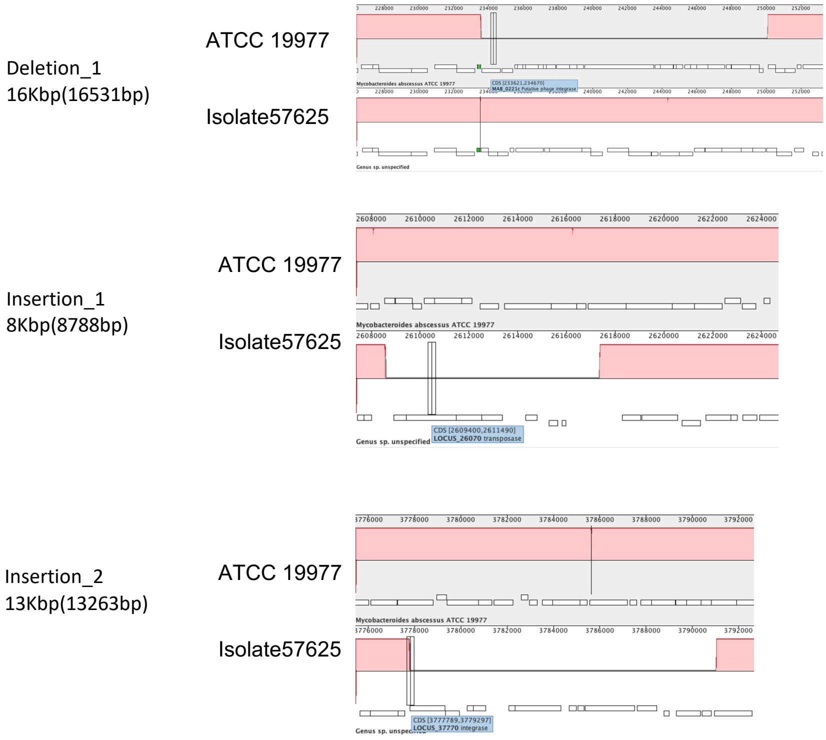


**Figure S3. The large SVs in Isolate57625**

One deletion and 2 insertions were detected in the isolate57625 chromosomes compared with ATCC19977. Phage integrase and transposase are present at the near region of these SVs.

A

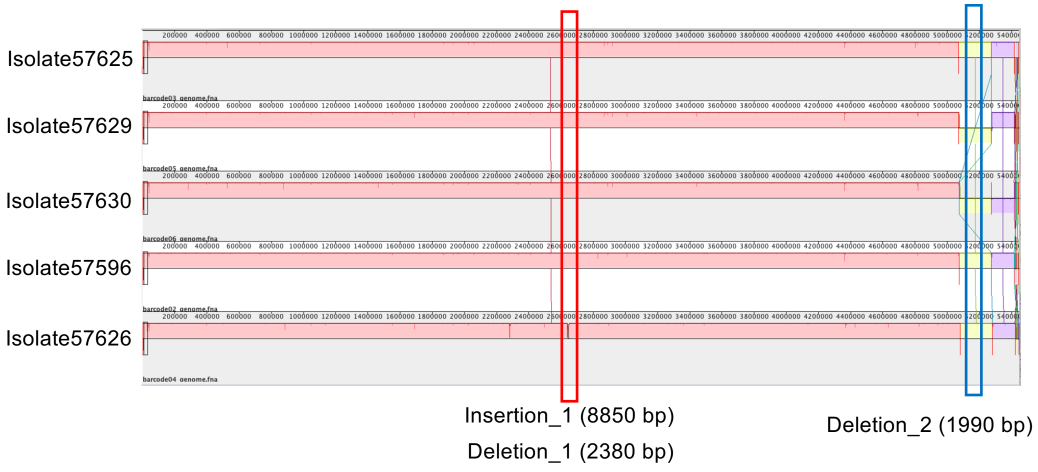


B

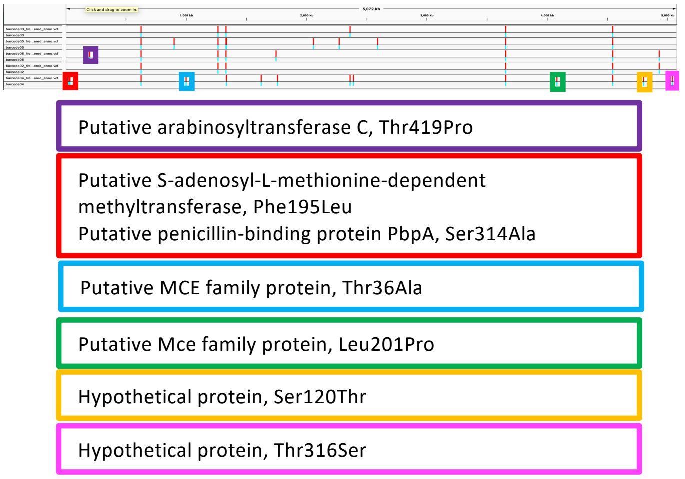


**Figure S4. The comparison of the genome of the isolates**

(A) One insertion and two deletions were detected in the Isolate57626 chromosome compared to the others.

(B) Six-missense mutation are present in the isolate genomes compares to that of Isolate57625.

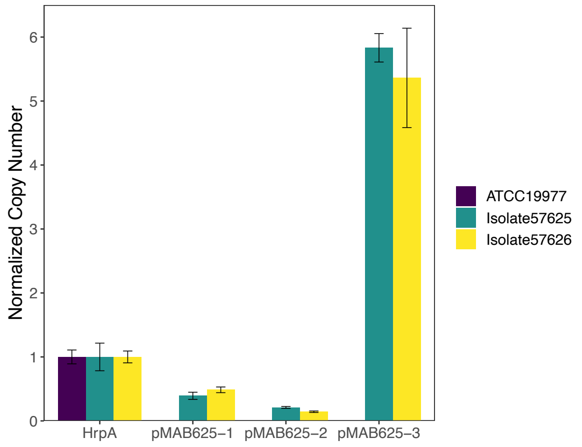


**Figure S5. pMAB625 plasmid copy number in the type strain and each isolate**

The copy number was analysed bu qPCR using primers specific to each plasmid. The normalised copy number represents the relative value that was normalized by the copy number of *HrpA* (encoded on the chromosome). The normalized copy number is the mean of three independent experiments and the error bar represents the standard deviation.

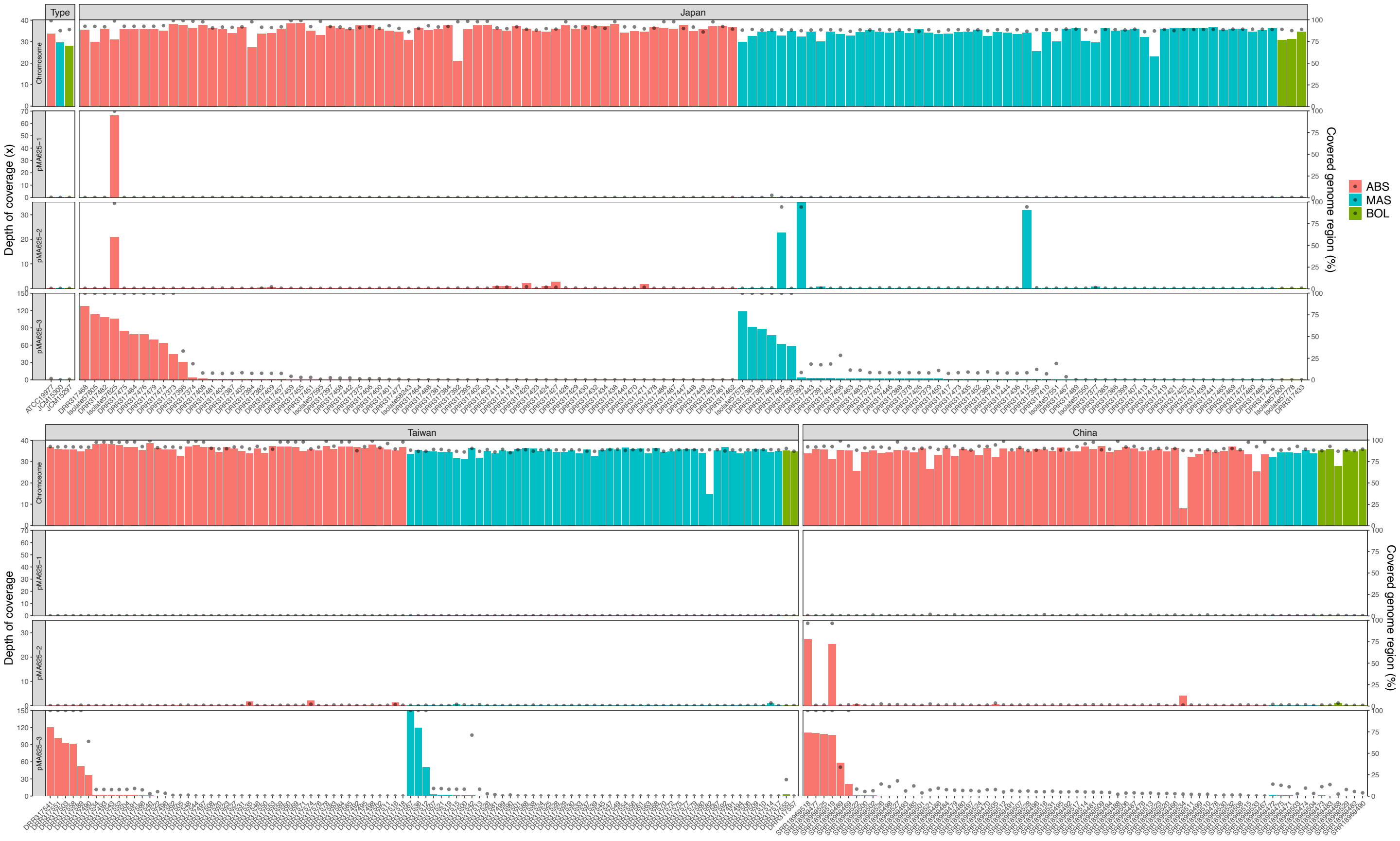


**Figure S6. The possession of pMAB625 plasmids in the isolates collected in East Asia**
Bars represent depth of coverage (x) and dots represent the covered genome region (%).
